# Longitudinal Auditory Pathophysiology Following Mild Blast Induced Trauma

**DOI:** 10.1101/2020.11.06.371591

**Authors:** Emily X. Han, Joseph M. Fernandez, Caitlin Swanberg, Riyi Shi, Edward L. Bartlett

## Abstract

Blast-induced hearing difficulties affect thousands of veterans and civilians. The long-term impact of even a mild blast exposure on the central auditory system is hypothesized to contribute to lasting behavioral complaints associated with mild blast traumatic brain injury (bTBI). Although recovery from mild blast has been studied separately over brief or long time windows, few, if any, studies have investigated recovery longitudinally over short-term and longer-term (months) time windows. Specifically, many peripheral measures of auditory function either recover or exhibit subclinical deficits, masking deficits in processing complex, real-world stimuli that may recover differently. Thus, examining the acute time course and pattern of neurophysiological impairment using appropriate stimuli is critical to better understanding and intervention of bTBI-induced auditory system impairments. Here, we compared auditory brainstem response, middle-latency auditory evoked potentials, and envelope following responses. Stimuli were clicks, tone pips, amplitude modulated tones in quiet and in noise, and speech-like stimuli (iterated rippled noise pitch contours) in adult male rats subjected to mild blast and sham exposure over the course of two months. We found that blast animals demonstrated drastic threshold increases and auditory transmission deficits immediately after blast exposure, followed by substantial recovery during the window of 7-14 days post-blast, though with some deficits remaining even after two months. Challenging conditions and speech-like stimuli can better elucidate mild bTBI-induced auditory deficit during this period. Our results suggest multiphasic recovery and therefore potentially different time windows for treatment, and deficits can be best observed using a small battery of sound stimuli.

**New and Noteworthy:** Few studies on blast-induced hearing deficits go beyond simple sounds and sparsely track post-exposure. Therefore, the recovery arc for potential therapies and real-world listening is poorly understood. Evidence suggested multiple recovery phases over 2 months post-exposure. Hearing thresholds largely recovered within 14 days and partially explained recovery. However, mid-latency responses, responses to AM in noise, and speech-like pitch sweeps exhibited extended changes, implying persistent central auditory deficits and the importance of subclinical threshold shifts.

## Introduction

Hearing loss stands out as one of the most commonly reported consequences following blast injuries and can last for months or even years without significant external injury (Cohen et al. 2002; Cave et al. 2007; Ritenour et al. 2008; Saunders et al. 2015). Most studies regarding blast-induced hearing loss have focused on damage in different parts of the peripheral auditory system (PAS) (Kerr 1980; DePalma et al. 2005), including hair cells, cochlear synapses, and auditory nerve damage. However, significant hearing difficulties can occur in the absence of peripheral diagnostic indicators such as eardrum rupture or clinical threshold shifts (hearing loss >25 dB), indicating potential disruptions upstream (Remenschneider et al. 2014; Saunders et al. 2015; Van Haesendonck et al. 2018).

Increasing clinical (Berger et al. 1997; Cohen et al. 2002; Cave et al. 2007; Ritenour et al. 2008; Lew et al. 2009; Gallun et al. 2012a) and laboratory (Patterson and Hamernik 1997; Ewert et al. 2012; Cho et al. 2013b; Du et al. 2013; Masri et al. 2018) findings suggest that the central auditory system (CAS) contains blast-susceptible structures. Subcortical CAS may be particularly vulnerable to blast injury, including mechanical damage and blood-brain barrier (BBB) permeability, excitotoxicity, and elevated markers of oxidative stress and neuroinflammation for at least 2 weeks (Knudsen and Øen 2003; Leung et al. 2008; Säljö et al. 2011; Cho et al. 2013a; Song et al. 2015; Walls et al. 2016). Functional changes, such as hyperactivity in the auditory brainstem (Luo et al. 2014a, 2014b) or structural changes in OHC loss (Ewert et al. 2012) or in the inferior colliculus (IC) and auditory thalamus (Mao et al. 2012), have been shown at 1-2 time points post-blast at various time points up to 2-3 weeks. Understanding the trajectory of post-blast recovery from primary and secondary damage can help to identify critical time points for diagnostics and therapies.

Clinical reports have suspected “hidden hearing loss” in blast-exposed veterans due to deficits in suprathreshold auditory processing with minimal changes in auditory thresholds (Gallun et al. 2012a; Saunders et al. 2015; Bressler et al. 2017) One consequence to this loss could be CAS adaptations to peripheral deafferentation (Caspary et al. 2005, 2008; Wang et al. 2009), which may lead to impaired temporal processing (Walton 2010; Parthasarathy and Bartlett 2011, 2012; Rabang et al. 2012). Blast studies on human subjects often used speech and complex temporally modulated stimuli to pin down “hidden” temporal processing losses at suprathreshold levels (Gallun et al. 2012b; Saunders et al. 2015; Bressler et al. 2017; Kubli et al. 2018). However, blast studies in animals rarely go beyond simple auditory stimuli (Ewert et al. 2012; Race et al. 2017; Masri et al. 2018).

In the current study, in addition to traditional measures, we chose Iterated Rippled Noise (IRN) to create a pitch contour with adjustable salience alongside Amplitude Modulation (AM) stimuli in quiet and in modulated noise as temporally complex stimuli in assessing the processing of temporal attributes. IRN has been used in neurophysiological and behavioral studies in both human (Krishnan et al. 2014, 2015; Peter et al. 2014; Thompson and Marozeau 2014; Wagner et al. 2017) and animal models (Bendor and Wang 2005; Alsindi et al. 2018).

## Materials and Methods

### Subject

Male Sprague-Dawley rats (3-4 months) were assigned into Sham group and Blast group randomly. A total of 11 Sham animals and 13 blast animals were used in this study. For a given sound stimulus, only complete sets of responses that included all time points were used for analysis. In a few sessions, there were recording sessions contaminated by movement artifact or movements that displaced electrode positions, and response sets affected by those were not included. All animals were kept and raised in relatively quiet and standard laboratory animal housing conditions. All protocols were approved by the Purdue Animals Care and Use Committee (PACUC #1111000280).

### Blast Exposure

Animals were anesthetized through intraperitoneal injection of a ketamine/xylazine cocktail (80 mg/kg and 10 mg/kg, respectively). The absence of eye-blink and paw-withdrawal reflexes was ensured prior to proceeding. Anesthetized animals were then placed on a platform beneath an open-ended shock tube to be exposed to the blast event, as described in our prior publications (Song et al. 2015; Walls et al. 2016; Race et al. 2017).

For the Blast group, each rat’s head was positioned beneath the open end of the shock tube such that the dorsum of the skull was the incident surface exposed to a composite blast (shock wave + blast wind). A custom plexiglass housing was temporarily placed over the animal’s torso for body protection to avoid cardiac or pulmonary effects of blast and to simulate the protective effects of military body armor (Rafaels et al. 2011). The head was fixed with a stereotaxic head frame with bite bar and ear bars (Kopf Instruments) to prevent blast wind-induced head acceleration. The blast exposure exhibited a recorded pressure profile with a rise to peak pressure within 0.3 msec, followed by overpressure and underpressure periods as follows: side-on (static) 150 kPa maximum overpressure, 1.25 msec overpressure duration, and 20 kPa minimum underpressure; face on (dynamic) 160 kPa maximum overpressure, 1.75 msec overpressure duration, and 5 kPa minimum underpressure. These conditions were the same as reported in our prior publications (Song et al. 2015; Walls et al. 2016; Race et al. 2017) and are considered to be a mild blast exposure, given the magnitude of the exposure and its single occurrence. All but one blast animal survived the exposure without displaying any motor or behavioral deficit during each animal’s longitudinal follow-up period.

Sham animals were placed equidistant from the blast source, but out of the path of the shockwave, therefore only exposed to the blast noise. Tympanic membrane integrity was verified for all Blast and Sham animals after injury using a surgical microscope.

### Auditory Evoked Potential Recordings

The animals underwent two-channel Auditory Evoked Potential (AEP) recordings at the following time points: pre-exposure (baseline), 1 day, 4 days, 7 days, 10 days, 14 days, 1 month, and 2 months. While the animals were under 1.8-2% isoflurane anesthesia, subdermal needle electrodes (Ambu) were inserted in the following locations (Fig. 1A): Channel 1 positive electrode was placed along the midline of the head (mid-sagittal) oriented Fz to Cz. Channel 2 positive electrode was positioned C3 to C4 along the interaural line. The negative/inverting electrode (used with positive electrodes for both channels 1 and 2) was placed under the mastoid of the right ear ipsilateral to the speaker. A ground electrode was placed in the back of the animal. These configurations were consistent with prior publications from our laboratory (Parthasarathy and Bartlett 2011, 2012; Parthasarathy et al. 2014; Lai and Bartlett 2015; Lai et al. 2017). Electrode impedances were confirmed to be less than 1 kΩ using a low impedance amplifier (RA4LI, TDT). After electrode placement, we subsequently sedated the animals by intramuscular injection of 0.2-0.3 mg/kg dexmedetomidine (Dexdomitor). AEP recordings were performed 10-15 min after removal from isoflurane to avoid anesthetic effects. The animals could respond to pain and acoustic stimuli but tend sit calmly under dexmedetomidine sedation, allowing about 3 hours of recording time.

**Fig 1.**
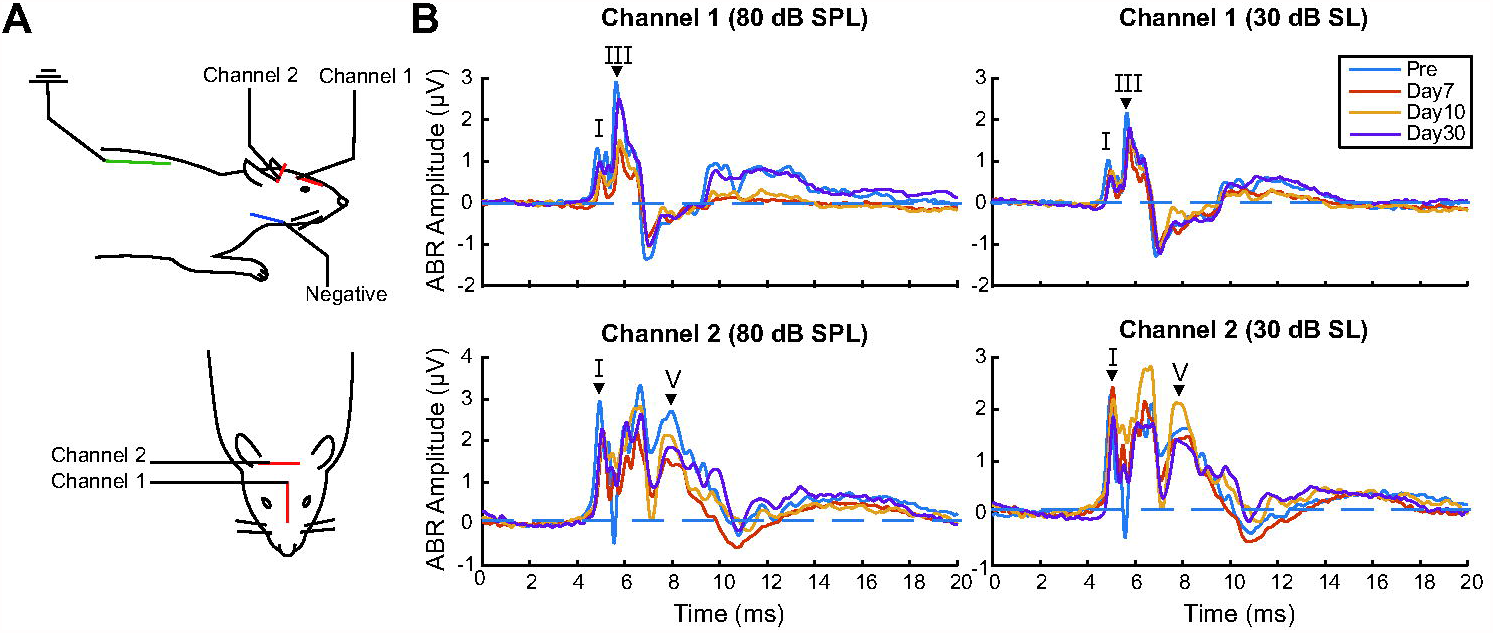
Auditory evoked potential experiment setup and examples of ABR waveforms. A) Electrode placement and channel configuration of the study’s auditory evoked potential experiment. B) Examples of ABR waveforms of a pre-exposure animal, at 80 dB SPL and 30 dB SL, with relevant wave peaks labeled. The waves for which amplitudes are measured are labeled with a black triangle.

Acoustic stimuli were presented free-field to the right ear (90□ azimuth) of animals, with directly in front of the animals’ face as the reference for 0□ azimuth, using a calibrated speaker (Bowers and Wilkins) at a distance of 115 cm directly facing the right ear. The measurements used in this study included auditory brainstem responses (ABRs), middle-latency responses (MLRs), envelop-following responses (EFRs) using AM in noise stimuli, and IRNs.

### ABR and MLR

6 Sham animals and 10 Blast animals were used in ABR analysis. For ABR, rectangular clicks (0.1 msec duration) and tone-pips (2 msec duration, 0.5 msec cos^2^ rise-fall time) with frequencies of 8 kHz and 16kHz were used. 8 kHz and 16 kHz were chosen based on previous findings: with 6-16 kHz being the most sensitive hearing region of rats, 8 kHz near the most sensitive region of normal rat audiogram (Parthasarathy et al. 2014) and hearing of frequencies higher than 8 kHz being most vulnerable to blast injury (Race et al. 2017). The sound levels of clicks and pips ranged from 90 to 10 dB peak SPL in 5-dB steps. All stimuli were presented in alternating polarity at 26.6 per second with 1500 repetitions (750 at each polarity). A 20 msec acquisition window (0-20 msec) was used.

Data were processed with a 30 Hz high-pass (HP) filter and a 3000 Hz low-pass (LP) filter prior to analysis. The ABR threshold was visually determined as the minimum sound level that produced a distinct ABR waveform, with confirmation from two other researchers. The ABR amplitudes of waves I and V from channel 2 were estimated as the differences of each wave’s amplitude, as seen in BioSigRP (TDT) and the baseline amplitude (measured as an average of 2 msec waveform prior to the cochlear microphonic).

6 Sham animals and 8 Blast animals were used in MLR analysis. For MLR, similar rectangular clicks and 8 kHz tone pips of alternating polarity as in ABR were used but were presented at a slower rate (3.33/sec vs. 26.6/sec in ABRs) and with a recording window of longer duration (100 msec vs. 20 msec in ABRs). This time window provides enough time to capture the stimulus-evoked “middle-latency” neural responses from the auditory midbrain, thalamus and cortex (Barth and Shi Di 1991; McGee et al. 1991; Di and Barth 1992; McGee and Kraus 1996; Phillips et al. 2011; Šuta et al. 2011) alongside ABR. Stimuli were presented at 80 dB sound pressure level (SPL) and 30 dB sensation level (SL, 30 dB above corresponding ABR thresholds), as determined in the previous ABR recordings. 1500 repetitions were collected over an acquisition time window of 100 msec to obtain an average response. Only one animal exhibited hearing threshold higher than 80 dB SPL at only one time point, for which MLR recording has been excluded for that point.

Channel 2 was used for MLR analyses, and results were qualitatively similar for channel 1. Data were processed with HP (fc = 10 Hz) and LP (fc = 300 Hz) filters prior to analysis.

### EFRs

EFRs were recorded during the same recording session following ABRs and MLRs using the same electrode configurations with similar techniques to Lai and Bartlett (2018) and Lai (Lai et al. 2017). The two channels were sensitive to a complementary range of amplitude modulation frequencies (AMFs) (Parthasarathy and Bartlett 2012), with channel 1 (mid-sagittal) being more sensitive to higher AMFs (90-2048 Hz) while channel 2 (interaural) is more sensitive to lower AMFs (8-90 Hz). The AM stimuli used for EFRs were sinusoidally amplitude-modulated (AM) sounds, with Gaussian noise, 8 kHz tone, or 16 kHz tone as carriers, and under 100% and 50% modulation depth with a stimulus duration of 200 msec. The AMFs selected for this study are 10 Hz, 45 Hz, and 256 Hz, based on the findings in Race at al. (Race et al. 2017), which found significant differences, particularly at the lower AMFs. The acquisition window was 300 msec long, and each response was an average of 200 repetitions. The stimuli were presented at 30 dB SL. For animals that had a hearing threshold above 70 dB SPL, which usually happens on day 1 post-exposure, EFR was not collected at the time point due to the limitation of the speaker and BiosigRP.

For AM in Noise stimuli, the same EFRs were used alongside a 71 Hz sinusoidally AM masker of the same length and onset, with Gaussian noise as the carrier, similar to Lai and Bartlett (Lai and Bartlett 2018). Noise AM maskers were presented at sound levels of 20dB SNR and 0SNR to the sound level of target AM. Prior to EFR amplitude analysis, data were passed through an LP filter of 3000 Hz and a high-pass filter that was either slightly below the AMF for AMFs <90 Hz, or 80 Hz for AMFs ≥ 90 Hz.

### IRNs

For 6 Sham animals and 8 Blast animals, IRNs were recorded during the same recording session following the previous stimuli using the same electrode configurations. The sound level of presentation was 30 dB SL (above click hearing threshold). Data for animals with a hearing threshold above 70 dB SPL were not collected at the time point.

IRN tone stimuli were created by sequential delay and add operations. Time-varying pitch curves were created by applying polynomial equations to create delays constructed from the fundamental frequencies of Chinese tone 2 and tone 4, delaying Gaussian noise (80 Hz-40 kHz) by the inversion of pitch and adding it back on itself in a recursive manner (Yost 1996a). The core MATLAB program used for generating IRN was modified from Krishnan et al. (Krishnan et al. 2014, 2015) This would generate dynamic, curvilinear pitch patterns (Swaminathan et al. 2008) that preserves variations in pitch using a broadband carrier. The number of iteration steps for these stimuli was 32, beyond which there is little or no change in pitch salience (Yost 1996b).

IRN iteration (ite) stimuli were created with the same polynomial equations used for tone 2, but with different iterations to create an array of IRN stimuli with different pitch salience. The numbers of iteration steps were 32, 16, 8, 4, and 2.

All IRN stimuli consisted of pairs of waveforms in original and inversed polarities to compensate for envelope or fine structure response under different calculations and cancel any microphonics. The stimulus duration was 250 msec, and the acquisition window was 300 msec long. Each response was an average of 200 repetitions. Given the main frequencies involved in the IRN autocorrelation (>100 Hz), channel 1 was used for IRN analyses, and results were qualitatively similar for channel 2.

### Statistics

Statistics were performed with statistics software JASP (Version 0.11, JASP Team, 2019). All statistics for ABR and EFR utilized 2-way repeated measures ANOVA test (α = 0.05) to check the significance of each main effect and interaction, undergoing Greenhouse-Geisser sphericity corrections (Greenhouse and Geisser 1959) and Tukey Post Hoc corrections (Tukey 1949). For ABR statistics, Wave I (channel 2), III (channel 1) and V (channel 2) were measured at each time point (Fig. 1B), corresponding to the auditory nerve (Wave I), cochlear nucleus (Wave III), and rostral brainstem/IC sources (Wave V) (Parthasarathy and Bartlett 2012; Simpson and Prendergast 2013). For EFR statistics, responses were analyzed from channel 2 for 10 Hz and 45 Hz, and from channel 1 for 256 Hz (Parthasarathy and Bartlett 2012). Prior to statistical tests, EFR amplitudes at signal frequencies were acquired through Fast Fourier Transformation (FFT) in MATLAB (MathWorks) similar to (Lai and Bartlett 2018).

For MLR statistics, P1, N1, P2, and N2 (Fig. 5A) peaks were measured at each time point, corresponding to subcortical (P1), thalamocortical (N1) and cortical sources (P2, N2) (Simpson and Prendergast 2013). Peak amplitudes were normalized to the pre-blast exposure baseline measurements for display in Fig. 5C, D. The normalized peak amplitudes at each time point were compared to the pre-stimulus baseline using a paired sign-rank test, with a 0.05 significance criterion.

For IRN statistics, we performed moving-window autocorrelations in 25 msec moving windows (5 msec steps) on each response waveform to simulate physiological tracking of temporal periodicity. Peak autocorrelation frequency was defined by the inverse of the time lag where peak autocorrelation value occurs in each window. This process yielded a peak frequency that reflect the frequency representation of the IRN auditory response for each of the 51 time windows in total (see Fig. 8B). Of those, 45 occurred during the stimulus. The peak frequencies were then compared to the “pseudopitches” of the IRN stimuli on corresponding time points. A value within 5 Hz of absolute difference to corresponding “pseudopitch” was considered “tracked.” We used this number of “tracked” peak frequencies, or “pitch-tracking score,” as a quantification for IRN performance. The significance of each main effect (time, blast condition, and IRN iterations) and interaction was assessed using similar 2-way repeated measures ANOVA test as ABR statistics (α = 0.05). For response-to-response correlation (Fig. 8D), the cross-correlation was measured between the response to the IRN stimuli pre-exposure and the response to the same stimulus post-exposure. Blast versus sham group was tested using the paired sign-rank test for this measure (α = 0.05).

## Results

### A. ABR and MLR

#### ABR Thresholds

Click ABR recordings captured distinctive courses of threshold changes over the two months post-exposure for blast and sham animals (Fig. 2). A large, >30dB SPL maximum threshold increase was observed in post-blast-exposure animals (Fig. 2, red lines). Adjacent animals exposed only to blast noise (Sham) did not undergo significant threshold shifts (Fig. 2, blue lines). Thresholds for blast group animals showed clear recovery during the first two weeks, with the largest changes occurring between 4 days – 10 days. Thresholds for blast-exposed animals remained significantly elevated (worse) than those of sham animals throughout the two months post-exposure that were measured (Simple Main Effects, day 30: df=1.000, F=10.904, p=0.005; day 60: df=1.000, F=12.727, p=0.003). Significant main effects of both Group (df=1.000, F=61.943, p=<0.001, η^2^_p_=0.816) and Time Point (df=2.554, F=41.932, p<0.001, η^2^_p_=0.750), as well as a significant Group*Time Point interaction effect (df=2.554, F=23.503, p<0.001, η^2^_p_=0.627), were observed.

**Fig 2.**
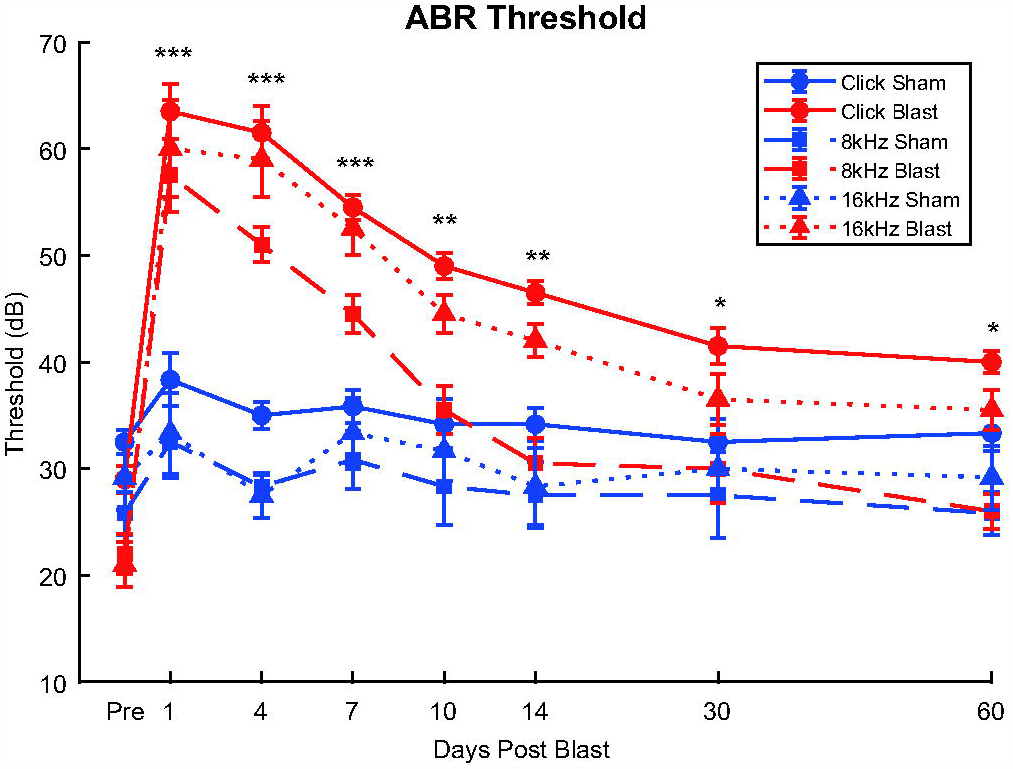
ABR threshold changes of Blast (N=10) and Sham (N=6) rats during the first two months post-exposure. Blast animals demonstrated drastic increases (worse) of Click, 8 kHz, and 16 kHz thresholds (red lines) post-exposure as opposed to Sham animals (blue lines). Significant main effects (p≤0.001) of Groups and Group*Time interactions were observed in all carriers. Significant Simple Main Effect of single time points observed in various carriers throughout the two months. For subsequent figures, red lines will denote blast-exposed animals, and blue lines will denote sham-exposed animals. Asterisks indicate time points where significant Simple Main Effects of Group was demonstrated (Supp. Table 1): ***Blast threshold significantly higher than Sham in Click, 8kHz, and 16kHz, p<0.05; **Blast threshold significantly higher only in Click and 16 kHz; *Blast threshold significantly higher only in Click.

Similar trends were observed with tone ABR recordings of 8 kHz and 16 kHz (Fig. 2), with a significant (p≤0.001) >30 dB increase in threshold within 48 hours post-blast-exposure and most prominent recovery between 4 days – 10 days. 8 kHz threshold differences between blast conditions became non-significant ((Simple Main Effects, df=1.000, F=3.151, p=0.098) at 10 days post-blast. At two weeks post-exposure, 16 kHz thresholds remained significantly elevated (Simple Main Effects, df=1.000, F=16.527, p<0.001), after which point the thresholds for the two chosen tone frequencies were no longer significantly different between Blast and Sham. Our rmANOVA analysis using Group and Time Points as factors showed significant main effects of Group (8 kHz: df=1.000, F=10.847, p=0.005, η^2^_p_=0.437; 16 kHz: df=1.000, F=19.697, p<0.001, η^2^_p_=0.585), Time (8 kHz: df=3.924, F=25.837, p<0.001, η^2^_p_=0.649; 16 kHz: df=3.043, F=20.181, p<0.001, η^2^_p_=0.590) and Group*Time Point interaction (8 kHz: df=3.924, F=13.490, p<0.001, η^2^_p_=0.491; 16 kHz: df=3.043, F=15.860, p<0.001, η^2^_p_=0.531) for 8 kHz and 16 kHz threshold respectively. These results demonstrate that broadband click thresholds remain significantly elevated over the 60 days measurement window. 8 kHz thresholds largely returned to baseline (Day 30: 8 dB difference, t=3.197, p=0.118; day 60: 4 dB difference, t=1.598, p=0.965) after two weeks, and 16 kHz thresholds remained significantly elevated compared to pre-blast baseline according to post hoc analysis (Day 30: 15.5 dB difference, t=5.687, p<0.001; day 60: 14.5 dB difference, t=5.320, p<0.001), although the difference between blast and Sham was not significant at these time points.

#### ABR Amplitudes

For our ABR and MLR measurements, we used two sound levels: 80 dB SPL was chosen because it is commonly used in auditory evoked potential studies in rat and human studies (Simpson et al. 1985; Alvarado et al. 2012; Race et al. 2017), and it elicits clear ABR responses in all except the most extreme cases of blast-exposure. In order to compensate for changes in threshold induced by blast exposure, we also measured ABR amplitudes at 30 dB SL above threshold (sensation level, or SL). This enabled us to separate changes in ABR amplitudes due to audibility (threshold) versus those due to threshold-independent changes in subcortical auditory signaling. Note that we did not attempt to compare later ABR waves with equivalent wave I amplitudes, as in Lai et al. (2017).

ABR wave amplitudes were assessed for wave I (putative auditory nerve), III (putative cochlear nuclei), and V (putative rostral brainstem and inferior colliculus) in response to click stimuli at 80 dB SPL (Fig. 3) and 30 dB SL (Fig. 4). Repeated measures statistics for 80 dB SPL and 30dB SL are shown in Tables 1-4.

**Fig 3.**
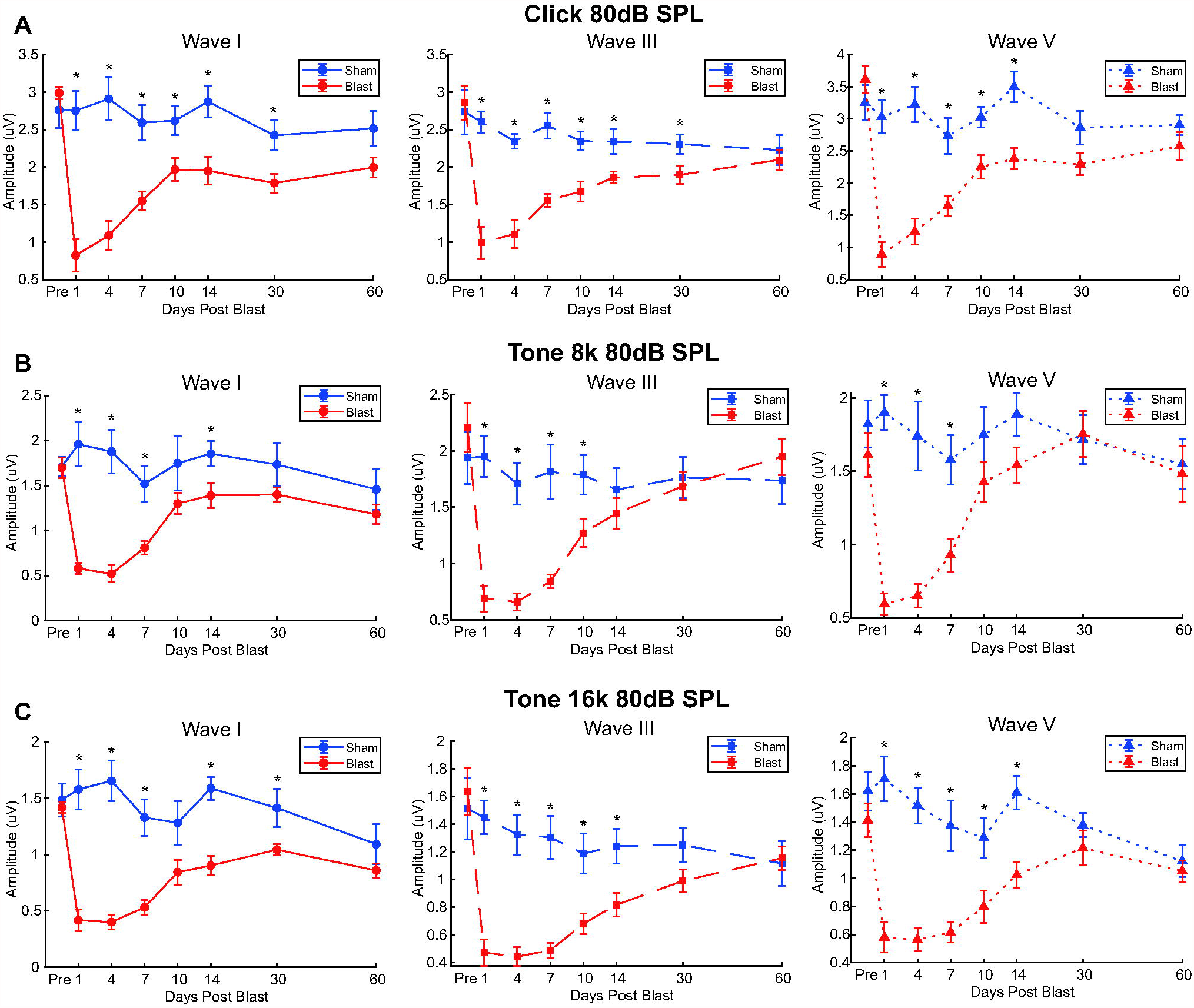
ABR wave I, III, and V amplitudes of Blast (N=10) and Sham (N=6) rats during the first two months post-exposure expose persistent blast-induced differences at 80 dB SPL. Significant main Group*Time interaction effects (p≤0.001) observed in waves I (left column), III (center column), and V (right column) for all carriers: A) Click ABR; B) 8 kHz ABR; C) 16 kHz ABR. Click ABR revealed blast-induced reduction of ABR wave amplitudes to a greater degree than both tone ABRs. Later waves (Wave III and V) showed earlier recovery in Blast animals. *Significant Simple Main Effect of Group in ABR Wave Amplitudes, p<0.05.

**Fig 4.**
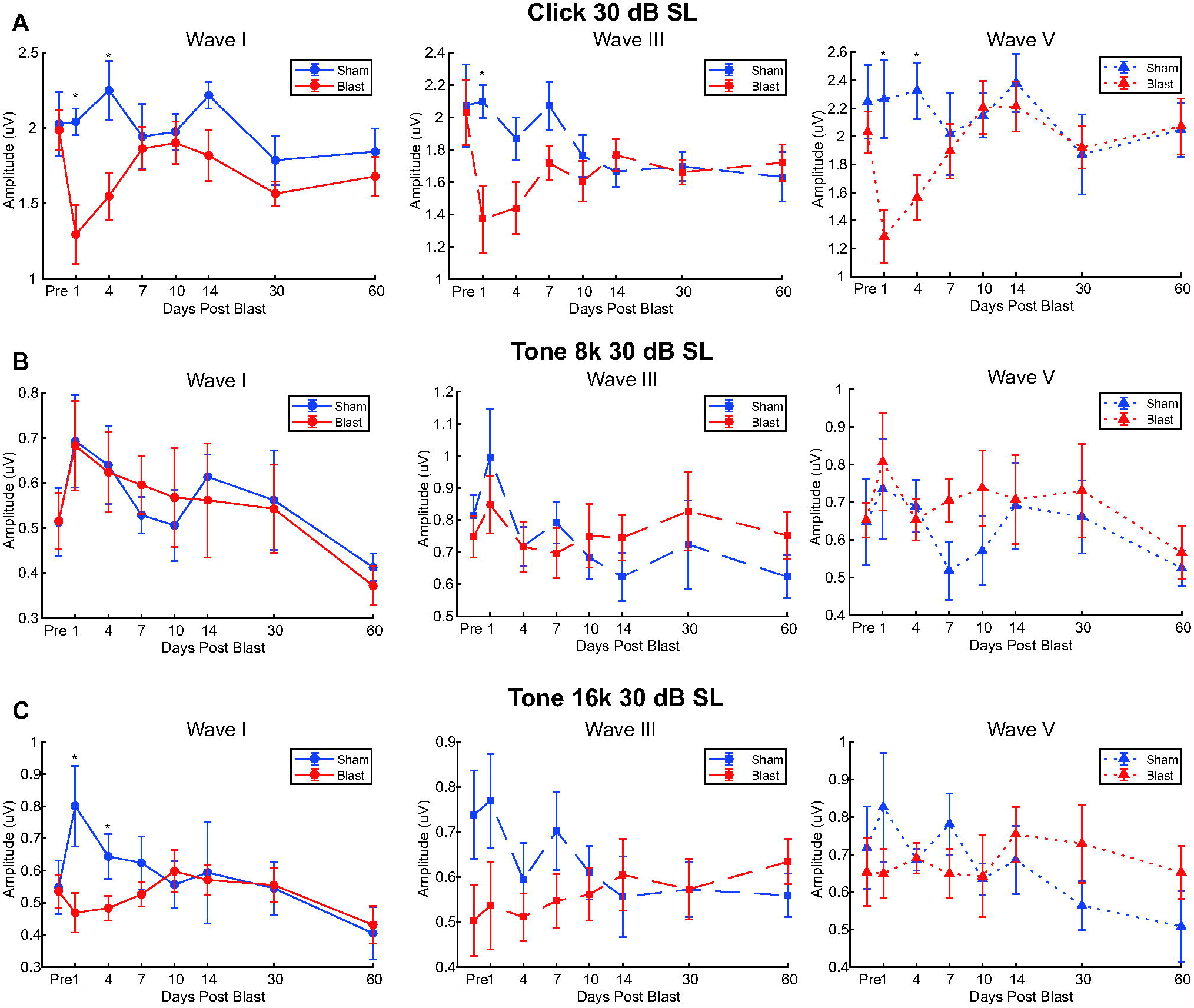
ABR wave I, III, and V amplitudes of Blast (N=10) and Sham (N=6) rats during the first two months post-exposure at 30 dB SL. Similar format to Fig. 3. Significant main Group*Time interaction effects only observed with Click ABR waves (Wave I: p=0.016; Wave III: p=0.04; Wave V: p=0.003) A) Click ABR; B) 8 kHz ABR; C) 16 kHz ABR. *Significant Simple Main Effect of Group in ABR Wave Amplitudes, p<0.05.

**Fig 5.**
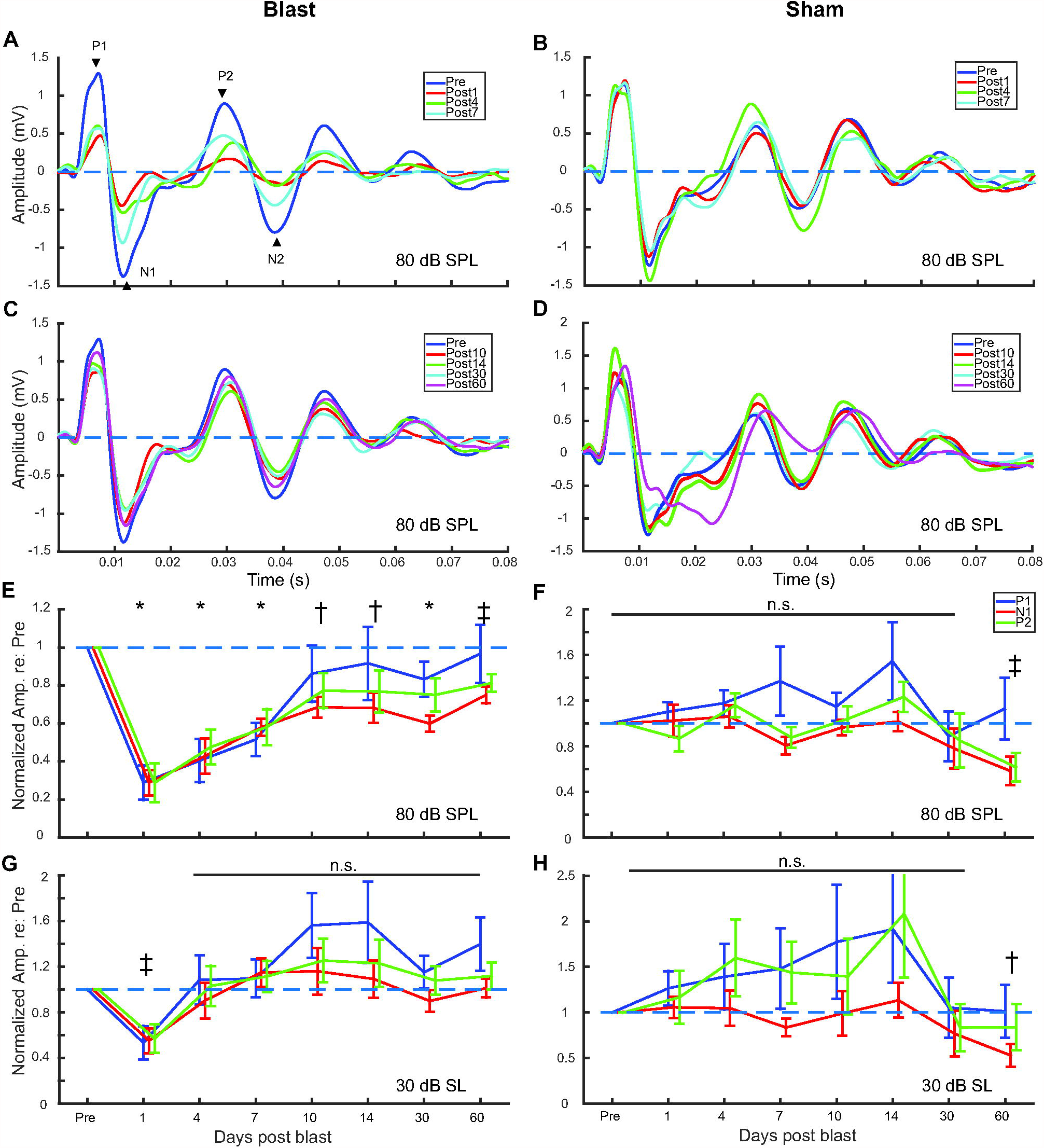
MLR waveforms and peak amplitudes of Blast (N=8) and Sham (N=6) rats during the first two months post-exposure at 80 dB SPL and at 30 dB SL (Thresh + 30 dB). Grand average traces of Click MLR waveforms at 80 dB SPL: A) Blast, pre-blast to day 7. Arrowheads indicate measured peaks in E-H; B) Sham, pre-blast to day 7; C) Blast, day 10 to day 60; D) Sham, day 10 to day 60. Normalized Click MLR wave amplitudes over time: E) Blast, 80 dB SPL; F) Sham, 80 dB SPL; G) Blast, 30 dB SL; H) Blast, 30 dB SL. *Significant difference in normalized wave P1, N1 and P2 amplitudes compared to pre-exposure, p<0.05. †Significant difference in normalized wave N1 amplitudes only, p<0.05. ‡Significant difference in normalized wave N1 and P2 amplitude, p<0.05.

**Table 1.**
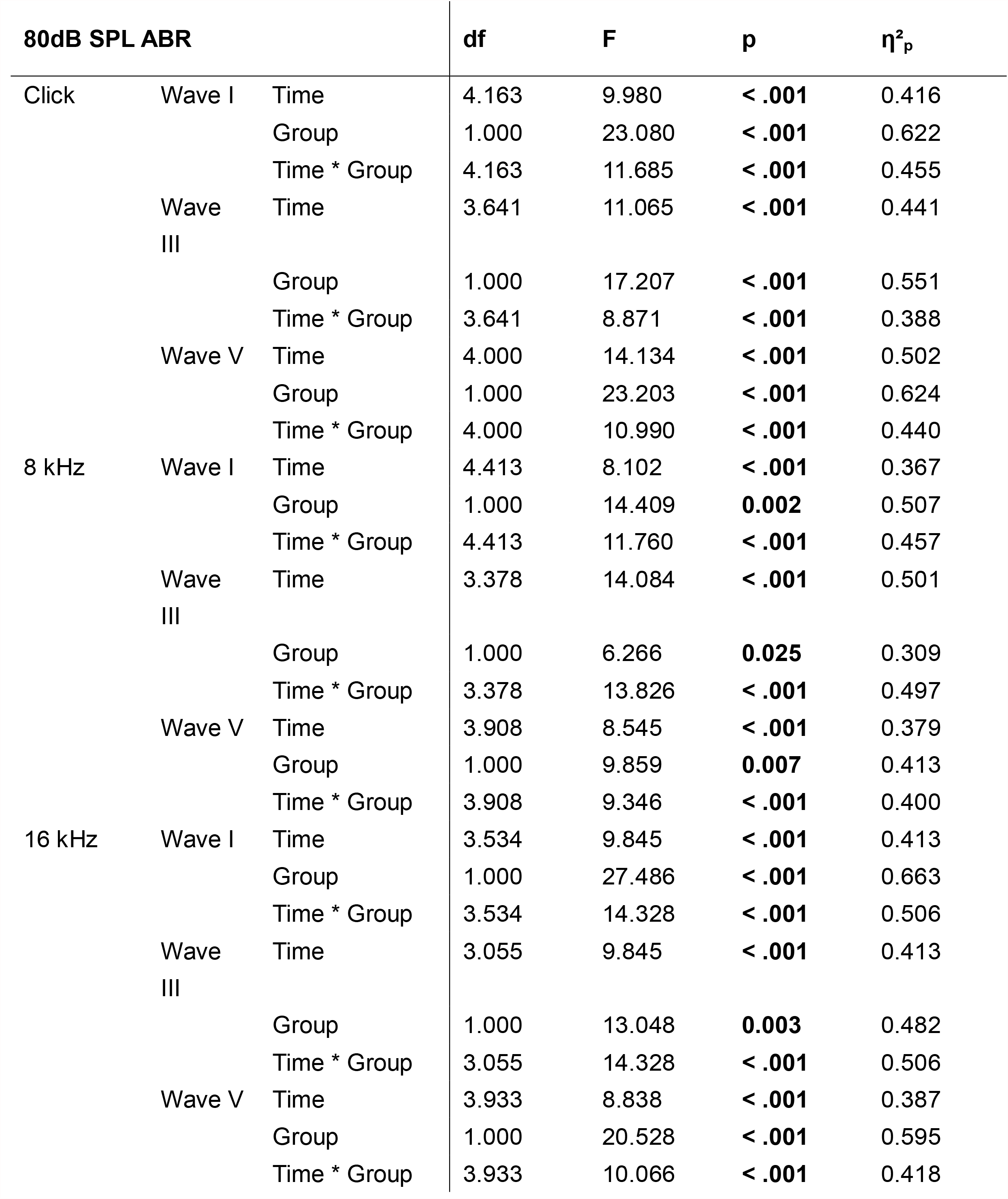
Summary of 80 dB SPL ABR Wave I, III and V repeated measure ABR statistics.

**Table 2.** Summary of 30 dB SL ABR Wave I, III and V repeated measure ABR statistics.

| 30dB SL ABR | | | df | F | p | $\eta^2_p$ |
| --- | --- | --- | --- | --- | --- | --- |
| Click | Wave I | Time | 4.360 | 2.554 | **0.043** | 0.154 |
|  |  | Group | 1.000 | 3.637 | 0.077 | 0.206 |
|  |  | Time * Group | 4.360 | 2.630 | **0.039** | 0.158 |
|  | Wave III | Time | 3.878 | 2.544 | 0.052 | 0.154 |
|  |  | Group | 1.000 | 1.811 | 0.200 | 0.115 |
|  |  | Time * Group | 3.878 | 2.720 | 0.040 | 0.163 |
|  | Wave V | Time | 4.784 | 2.568 | **0.037** | 0.155 |
|  |  | Group | 1.000 | 1.479 | 0.244 | 0.096 |
|  |  | Time * Group | 4.784 | 3.429 | **0.009** | 0.197 |
| 8 kHz | Wave I | Time | 4.264 | 3.146 | **0.018** | 0.183 |
|  |  | Group | 1.000 | 6.852e<sup>-5</sup> | 0.994 | 0.000 |
|  |  | Time * Group | 4.264 | 0.200 | 0.945 | 0.014 |
|  | Wave III | Time | 3.414 | 2.432 | 0.069 | 0.148 |
|  |  | Group | 1.000 | 0.020 | 0.889 | 0.001 |
|  |  | Time * Group | 3.414 | 1.184 | 0.328 | 0.078 |
|  | Wave V | Time | 3.912 | 1.837 | 0.136 | 0.116 |
|  |  | Group | 1.000 | 0.361 | 0.558 | 0.025 |
|  |  | Time * Group | 3.912 | 0.618 | 0.648 | 0.042 |
| 16 kHz | Wave I | Time | 4.469 | 2.325 | **0.031** | 0.142 |
|  |  | Group | 1.000 | 1.046 | 0.324 | 0.070 |
|  |  | Time * Group | 4.469 | 2.381 | **0.027** | 0.145 |
|  | Wave III | Time | 3.160 | 0.562 | 0.651 | 0.039 |
|  |  | Group | 1.000 | 1.149 | 0.302 | 0.076 |
|  |  | Time * Group | 3.160 | 1.930 | 0.136 | 0.121 |
|  | Wave V | Time | 4.874 | 1.160 | 0.338 | 0.077 |
|  |  | Group | 1.000 | 4.905e<sup>-4</sup> | 0.983 | 0.000 |
|  |  | Time * Group | 4.874 | 1.646 | 0.161 | 0.105 |

**Table 3.**
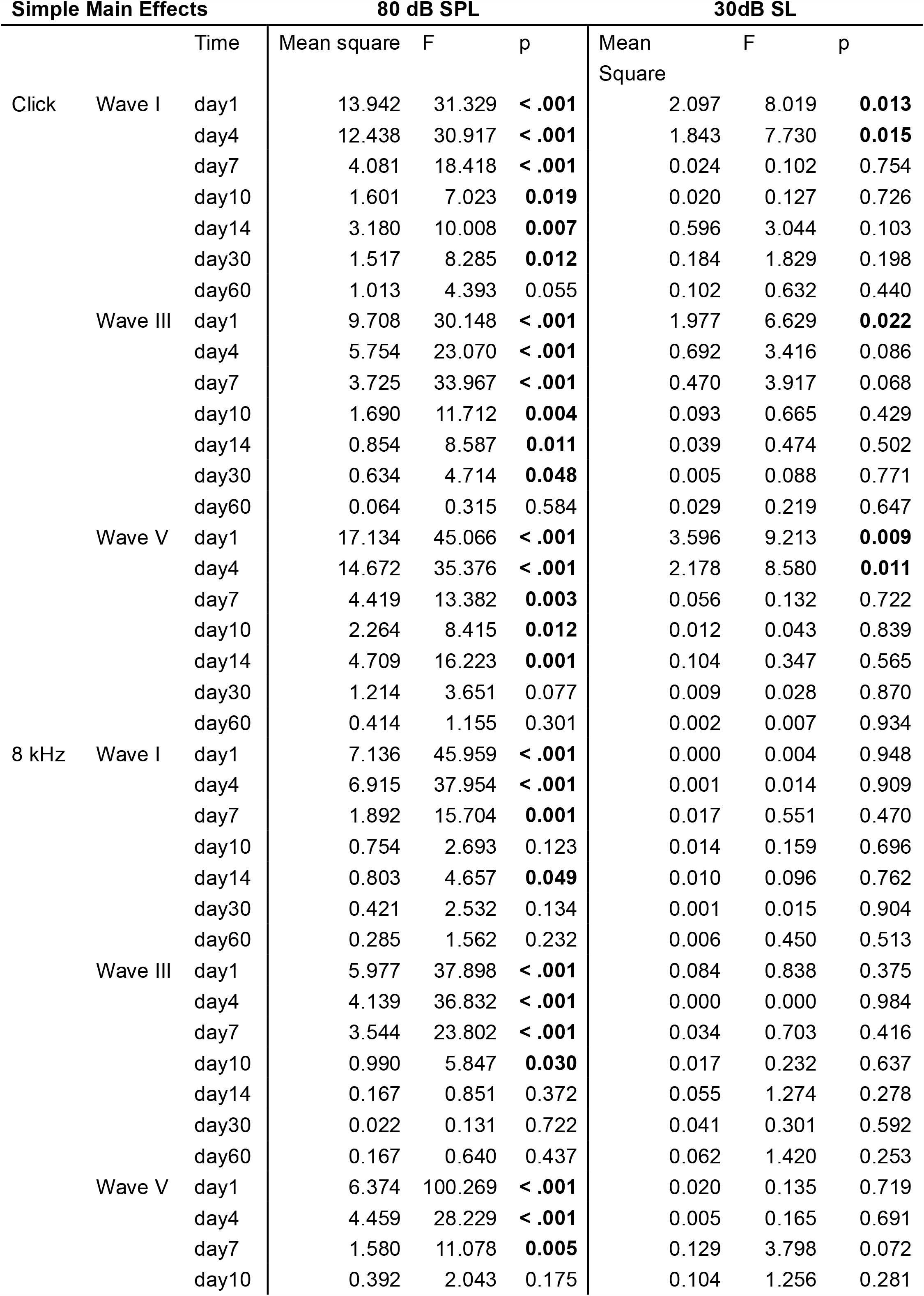
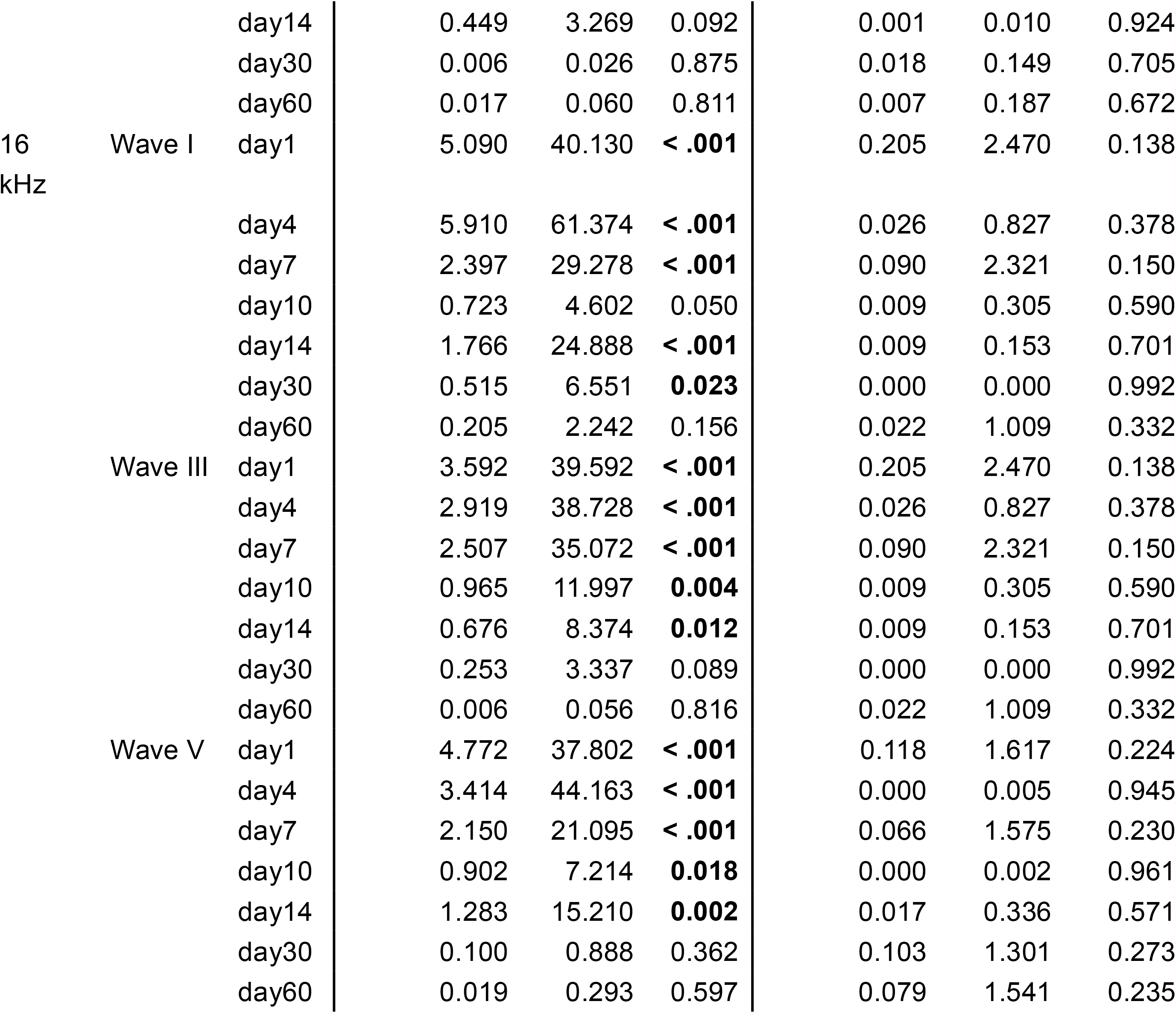
Simple main effects of Group on ABR wave amplitudes over time. Post-blast ABR amplitudes of Blast (N=10) and Sham (N=6) groups are compared using rmANOVA at each time point recorded. A p<0.05 showed significant simple main effect of Group at that time point.

**Table 4.**
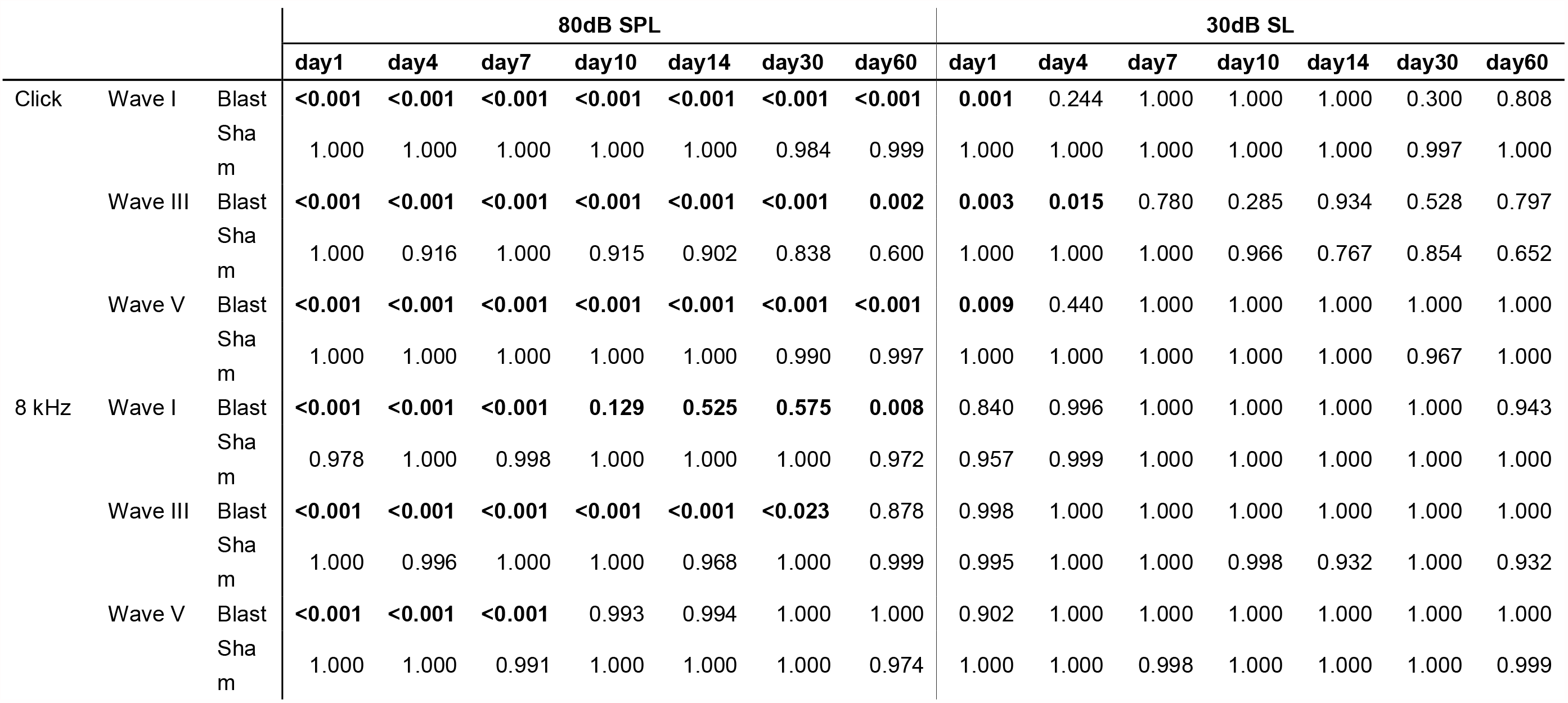
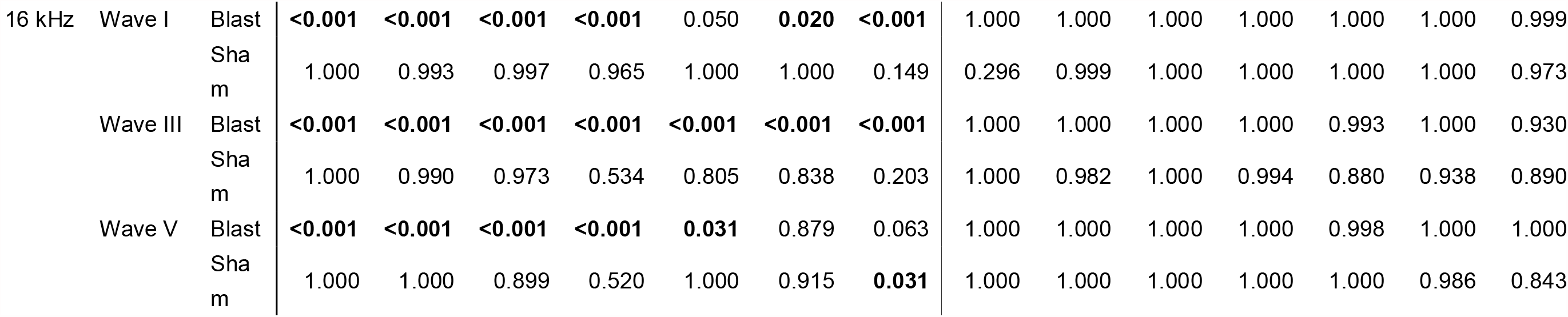
Summary of post hoc tests against pre-blast ABR amplitudes. Post-blast ABR amplitudes of Blast (N=10) and Sham (N=6) are compared against pre-blast amplitudes of the same group to show blast impact and recovery.

**Wave I:** Wave I amplitudes at 80 dB SPL for all ABR carriers at 80 dB SPL exhibited significant main effects of Group, Time, and Group*Time interaction (Table 1). Compared to pre-exposure responses, wave I amplitudes were significantly smaller at all time points tested in blast animals for clicks, 8 kHz tones, and 16 kHz tones, indicating lasting cochlear/auditory nerve damage (Table 4). No significant changes in wave I amplitudes were observed in Sham exposed animals at any time point.

**Wave III:** Wave III amplitudes at 80 dB SPL for all ABR carriers at 80 dB SPL exhibited significant main effects of Group, Time, and Group*Time interaction (Table 1), with Group effects lasting for 14 days for Click and 16 kHz tones and 10 days for 8 kHz tones. Compared to pre-exposure responses, wave III amplitudes were significantly smaller at all time points tested in blast animals for clicks and 16 kHz tones and up to 30 days for 8 kHz tones, indicating lasting declines in cochlear nucleus excitation (Table 4). No significant changes in wave III amplitudes were observed in Sham exposed animals at any time point.

**Wave V:** Wave V amplitudes at 80 dB SPL for all ABR carriers at 80 dB SPL exhibited significant main effects of Group, Time, and Group*Time interaction (Table 1), with Group effects lasting for 14 days for Click and 16 kHz tones and 7 days for 8 kHz tones. Compared to pre-exposure responses, wave V amplitudes were significantly smaller at all time points tested in blast animals for clicks, indicating lasting declines in rostral brainstem/IC excitation for brief, broadband clicks (Table 4). However, decreases in wave V amplitudes persisted for only 7 days for 8 kHz tones and 14 days for 16 kHz tones, suggesting that despite decreases in cochlear nucleus excitation (as represented by wave III amplitude), rostral brainstem/IC responses compensated and restored their responses. No significant changes in wave V amplitudes were observed in Sham exposed animals at any time point except for a small decline for 16 kHz responses 60 days post Sham exposure (Table 4).

The effects on ABR waves were greatly diminished when responses to 30 dB SL sounds were measured, as shown in Table 2 and Table 3. For Wave I, significant main effects of Time (Click: df=4.360, F=2.554, p=0.043, η^2^_p_=0.154; 8 kHz: df=4.264, F=3.146, p=0.018, η^2^_p_=0.183; 16kHz: df=4.469, F=2.325, p=0.031, η^2^_p_=0.142) but not Group (Click: df=1.000, F=3.637, p=0.077, η^2^_p_=0.206; 8 kHz: df=1.000, F<0.001, p=0.994, η^2^_p_<0.001; 16kHz: df=1.000, F=1.046, p=0.324, η^2^_p_=0.070) were observed for click, 8 kHz, and 16 kHz. Additionally, significant Group*Time interaction effects were only observed for Click (df=4.360, F=2.630, p=0.039, η^2^_p_=0.158) and 16 kHz (df=4.469, F=2.381, p=0.027, η^2^_p_=0.145). Simple main effects of Group (df=1.000) were only observed in Click (Table 3).

Compared to pre-exposure responses, wave I and V responses to clicks were significantly reduced 1 day post-blast and wave III responses were significantly reduced days 1-4. Otherwise, there were no significant declines in wave amplitudes in the Blast group, and there were no significant amplitude changes in the Sham group.

### B. MLR

In order to observe thalamocortical and cortical neural transmission in response to acoustic transients, we recorded middle-latency auditory responses to click and 8 kHz tone stimuli. These stimuli were identical to those used for ABR, but the presentation rate was much slower, and the analysis window and filters were different (see Methods). Measurements were made for the first four main peaks of the MLR. Here, P1 corresponds to subcortical activity, largely encompassing the ABR. N1 corresponds to thalamocortical transmission, while P2 and N2 are thought to correspond to primarily cortical activity (Deiber et al. 1988; Liégeois-Chauvel et al. 1994; Tichko and Skoe 2017; Musiek and Nagle 2018).

#### 80 dB SPL responses

In blast animals, all waves were decreased relative to pre-blast baseline for days 1-7 post-blast (p<0.05, sign-rank test) in response to 80 dB SPL click stimuli. Simple Main effect of blast showed similar results for P1, N1 and P2 (Table 5). Grand average traces are shown for MLR responses in this time window in Fig. 5A, relative to the pre-blast waveform (thick blue line in A-D). Even after the blast, the morphology and timing of the MLR waveform remained relatively intact, but the amplitudes were significantly diminished, shown as a significant Time*Group interaction effect for all three waves of interest (P1: df=2.942, F=4.111, p=0.014, η^2^_p_=0.255; N1: df=3.460, F=7.786, p<0.001, η^2^_p_=0.393; P2: df=3.684, F=5.607, p=0.001, η^2^_p_=0.318). In Fig. 5E-H, wave amplitudes were normalized to the pre-blast waves and measured.

**Table 5.**
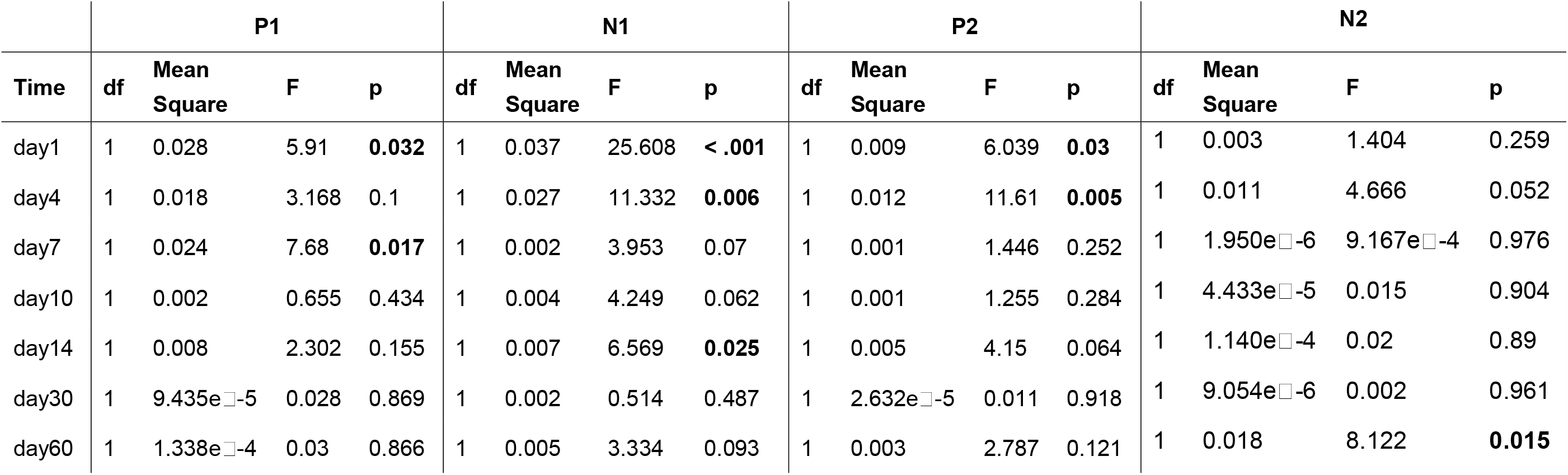
Simple main effects of Group on click MLR wave amplitudes at 80 dB SPL over time. Post-blast click MLR amplitudes of Blast (N=8) and Sham (N=6) groups at 80 dB SPL are compared using rmANOVA at each time point recorded. A p<0.05 showed significant simple main effect of Group at that time point.

|  | P1 |  |  |  | N1 |  |  |  | P2 |  |  |  | N2 |  |  |  |
| --- | --- | --- | --- | --- | --- | --- | --- | --- | --- | --- | --- | --- | --- | --- | --- | --- |
| Time | df | Mean Square | F | p | df | Mean Square | F | p | df | Mean Square | F | p | df | Mean Square | F | p |
| day1 | 1 | 0.028 | 5.91 | **0.032** | 1 | 0.037 | 25.608 | **< .001** | 1 | 0.009 | 6.039 | **0.03** | 1 | 0.003 | 1.404 | 0.259 |
| day4 | 1 | 0.018 | 3.168 | 0.1 | 1 | 0.027 | 11.332 | **0.006** | 1 | 0.012 | 11.61 | **0.005** | 1 | 0.011 | 4.666 | 0.052 |
| day7 | 1 | 0.024 | 7.68 | **0.017** | 1 | 0.002 | 3.953 | 0.07 | 1 | 0.001 | 1.446 | 0.252 | 1 | 1.950e<sup>-6</sup> | 9.167e<sup>-4</sup> | 0.976 |
| day10 | 1 | 0.002 | 0.655 | 0.434 | 1 | 0.004 | 4.249 | 0.062 | 1 | 0.001 | 1.255 | 0.284 | 1 | 4.433e<sup>-5</sup> | 0.015 | 0.904 |
| day14 | 1 | 0.008 | 2.302 | 0.155 | 1 | 0.007 | 6.569 | **0.025** | 1 | 0.005 | 4.15 | 0.064 | 1 | 1.140e<sup>-4</sup> | 0.02 | 0.89 |
| day30 | 1 | 9.435e<sup>-5</sup> | 0.028 | 0.869 | 1 | 0.002 | 0.514 | 0.487 | 1 | 2.632e<sup>-5</sup> | 0.011 | 0.918 | 1 | 9.054e<sup>-6</sup> | 0.002 | 0.961 |
| day60 | 1 | 1.338e<sup>-4</sup> | 0.03 | 0.866 | 1 | 0.005 | 3.334 | 0.093 | 1 | 0.003 | 2.787 | 0.121 | 1 | 0.018 | 8.122 | **0.015** |

Between 7 and 10 days, the early P1 wave recovers to within 10-15% of its baseline amplitude, whereas the later waves recovered more slowly (Fig. 5E). In particular, the N1 wave, thought to reflect thalamocortical transmission (Barth and Shi Di 1991; McGee et al. 1991, 1992; Di and Barth 1992; Brett et al. 1996; McGee and Kraus 1996; Phillips et al. 2011; Šuta et al. 2011), remained significantly lower in blast animals even 60 days post-blast (p<0.05, sign-rank test, Fig. 5E). By contrast, the MLR waves in sham animals were largely stable across the measurement time (Fig. 5F). Although there was some decline in the later waves for the last time window, this was not statistically significant (Fig. 5B, D, F).

MLR responses to 8 kHz, 80 dB SPL tone pips largely mirrored the results to clicks, with significant decreases for all waves for post-blast days 1-7 and a lasting decline in N1 for the duration of measurements (p<0.05, sign-rank test, traces not shown). Sham responses did not show any significant changes in MLR waves in response to the 80 dB SPL tone pips.

#### 30 dB SL

MLR responses to clicks at 30 dB SL were reduced in Blast animals 1 day after the blast but recovered to baseline levels afterwards. There was a tendency towards elevated P1 amplitudes, but this was not significant (Fig. 5G). Sham animals did not show any significant changes, though there was a tendency towards an increase in wave amplitude (Fig. 5H). Similar results were found for responses to tones at 30 dB SL (not shown).

### C. EFR and EFR in noise

Given the different time courses and extents of ABR threshold change for clicks and tones, we measured the corresponding EFRs in response to Gaussian broadband noise (nSAM), 8 kHz, and 16 kHz sinusoidal tone carriers. Considering that slow AM (<50 Hz) and faster AM (>50 Hz) are differentially represented throughout cortical and subcortical auditory nuclei (Joris et al. 2004; Wang et al. 2008), three representative AMFs (10, 45, and 256 Hz) were selected from previous publications (Parthasarathy et al. 2010, 2014; Parthasarathy and Bartlett 2011, 2012; Race et al. 2017) and tested in quiet at 100% and 50% modulation depth. AM stimuli were also presented at 30 dB SL with a 71 Hz sinusoidally AM masker of the same length and onset, with Gaussian noise as the carrier, at 20dB SNR and 0 SNR relative to the sound level of target AM. Responses were collected from both electrodes, but for 10 and 45 Hz AMFs, channel 2 responses were analyzed; and for 256 Hz AMF, channel 1 responses were analyzed (see Methods). For each carrier, simple main effects of all conditions were analyzed.

#### EFRs in quiet

For all three carriers in quiet, EFR amplitudes were similar at 10 and 256 Hz across time points and AM modulation depths (Fig. 6). Overall, the nSAM FFT amplitudes were higher in the Blast group in quiet (df=5.000, F=9.629, p=0.008, η^2^_p_=0.426), with 45 Hz being the most affected. Interestingly, in contrast to the lower FFT Amplitude found in Blast AM at 80 dB SPL (Race et al. 2017), when 30 dB sensation level (threshold +30 dB) was to compensate for threshold differences, FFT amplitude of 45 Hz nSAM was higher in Blast than in Sham animals (Fig. 6B). This difference was most salient on day 7 for 45 Hz nSAM (Post hoc comparison: t=-4.122, p=0.006). For 8 kHz SAM and 16 kHz SAM, the slight elevation of AM FFT Amplitude in Blast animals was not significant (Fig. 6C and 6D). Surprisingly, time did not have a significant interaction across repeated measures for AM response with any carrier either.

**Fig 6.**
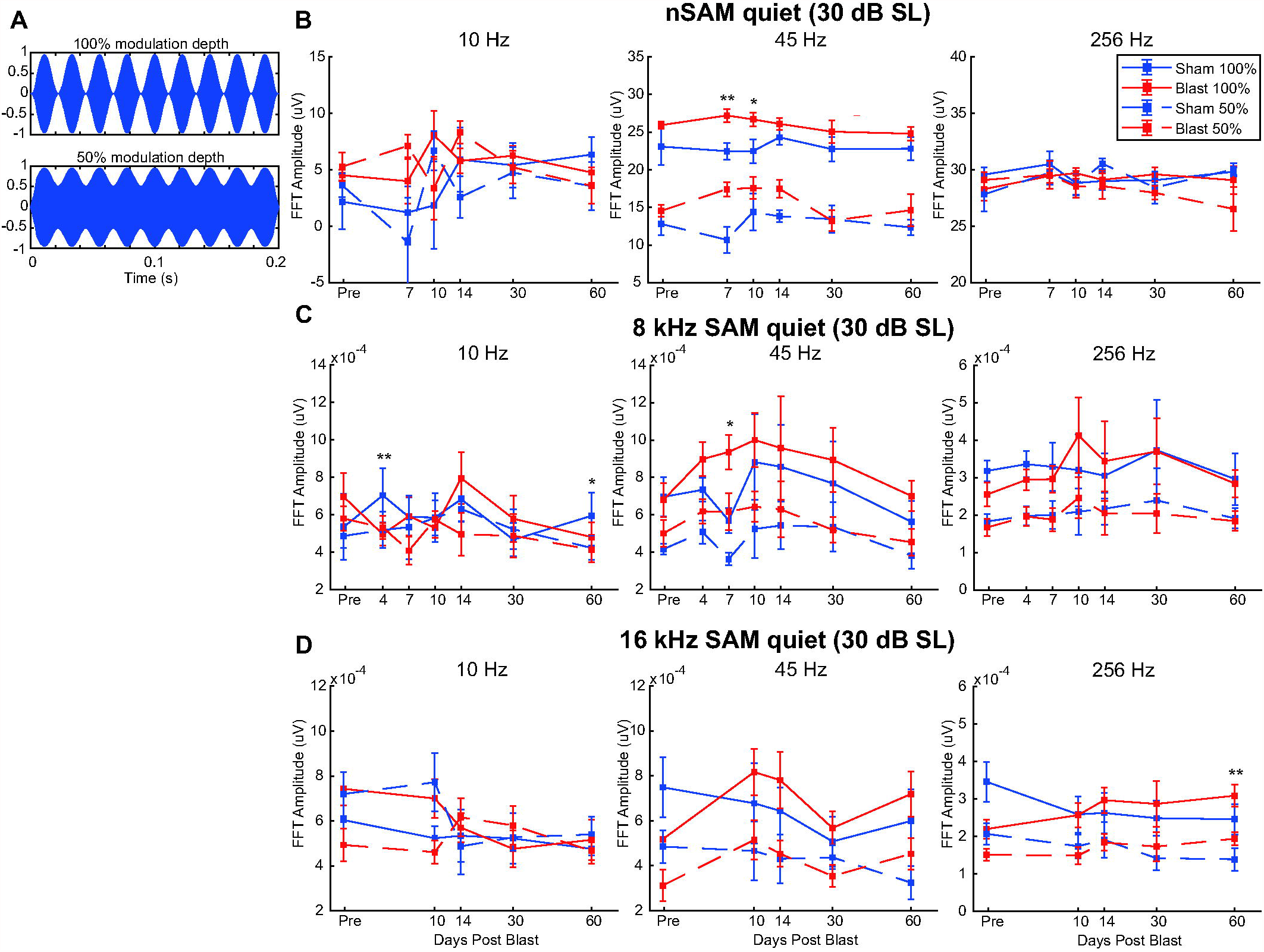
AM depth stimuli and EFR responses from Blast (N=10) and Sham (N=6) rats during the first two months post-exposure at 30 dB above threshold, in quiet. A) AM depth stimulus waveforms at 100% and 50% modulation depths; B) nSAM FFT amplitudes at 10 Hz (left), 45 Hz (center), and 256 Hz (right). Significant Group effect at 45Hz (p=0.007); Similar format in C and D. C) SAM 8 kHz FFT amplitudes at 45 Hz show a steady yet insignificant increase in later short-term (day 7-14); D) SAM 16k FFT amplitudes found no significant Group effect. **Significant Simple Main Effect of Group in FFT Amplitudes in both 100% depth and 50% depth *Significant Simple Main Effect of Group in FFT Amplitudes only in 100% depth

#### EFR in noise

Not surprisingly, Noise level and Modulation Depth both had a significant repeated measures effect on nSAM (Noise level: df=2.000, F=263.217, p<0.001, η^2^_p_=0.953; Depth: df=1.000, F=455.655, p<0.001, η^2^_p_=0.972), 8 kHz SAM (Noise level: df=2.000, F=19.308, p<0.001, η^2^_p_=0.580; Depth: df=1.000, F=72.031, p<0.001, η^2^_p_=0.837) and 16 kHz SAM (Noise level: df=2.000, F=16.691, p<0.001, η^2^_p_=0.544; Depth: df=1.000, F=49.742, p<0.001, η^2^_p_=0.780). Noise level and Depth also have a significant interaction effect with Groups for nSAM overall (Noise level: df=2.000, F=10.295, p<0.001, η^2^_p_=0.442; Depth: df=1.000, F=6.057, p=0.029, η^2^_p_=0.318), showing blast nSAM responses as less affected 20 SNR noise, but more sensitive to AM attenuation for lower modulation depth (Fig. 7B). Noise level also affect sham animals less than blast animals for 8 kHz SAM overall, showing a significant interaction effect with Group (df=2.000, F=5.696, p=0.008, η^2^_p_=0.289, Fig. 7C). These conditions do not have significant interaction effects with Group on 16 kHz SAM (data not shown). Noise level had significant interaction effects with both Group (df=2.000, F=6.130, p=0.011, η^2^_p_=0.320) and Depth (df=2.000, F=19.438, p <0.001, η^2^_p_=0.599) for nSAM 45 Hz, while the effect of Time or Depth between Groups is not significantly different for any modulation frequency.

**Fig 7.**
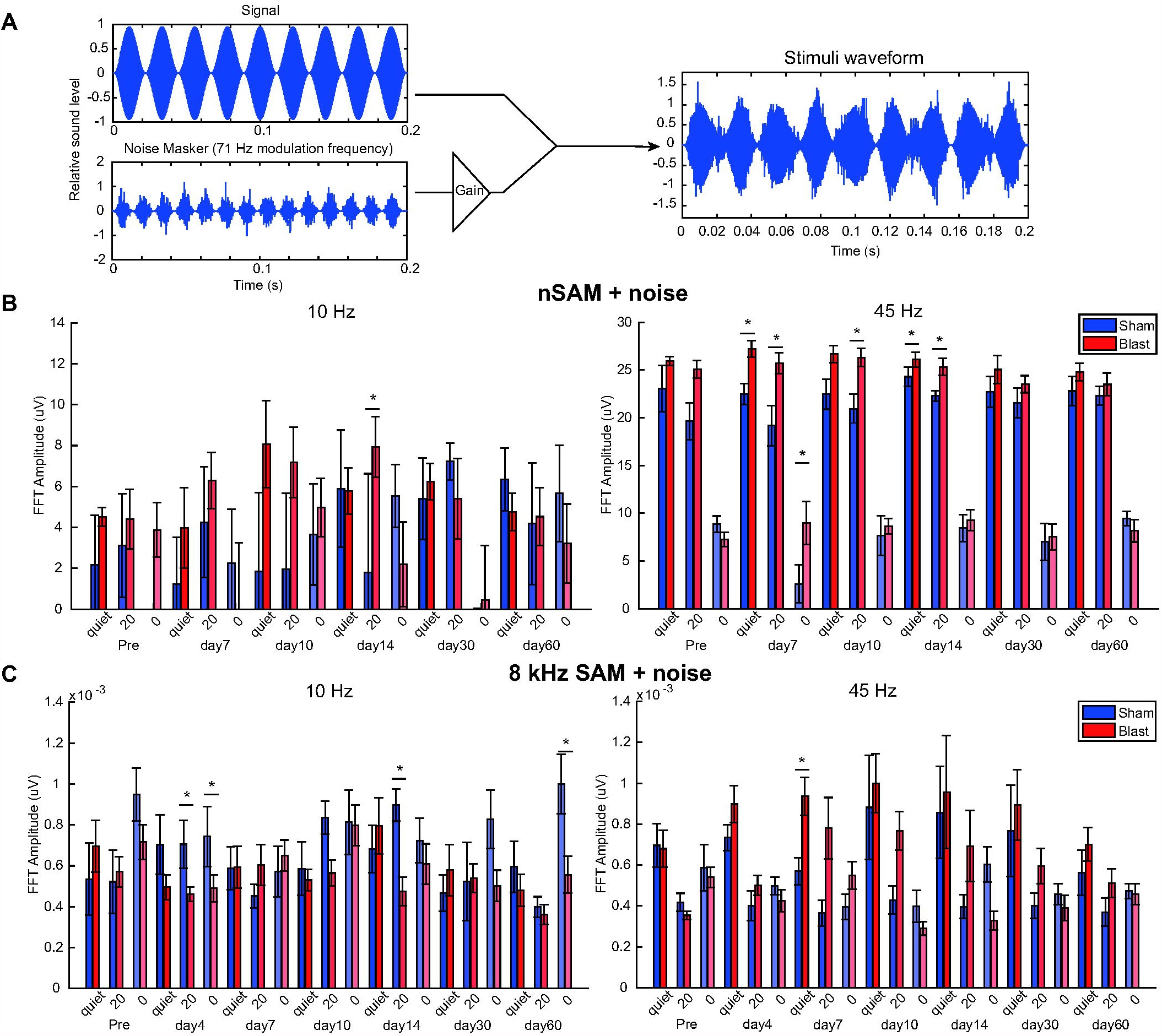
AM noise stimuli and EFR responses from Blast (N=10) and Sham (N=6) rats during the first two months post-exposure at 30 dB above threshold, modulation depth 100%. A) AM noise stimulus composition and waveform. B) Amplitude modulated noise carrier. FFT amplitudes at signal modulation frequency in quiet and with 71 Hz AM noise masker level of 20SNR or 0SNR (equal) show significant Noise * Group effect for: B) nSAM noise at 45 Hz (p=0.011) modulation frequency; C) SAM 8 kHz noise at 10 Hz (p=0.001) and 45 Hz (p=0.015) modulation frequency. * Significant Simple Main Effect of Group

**Fig 8.**
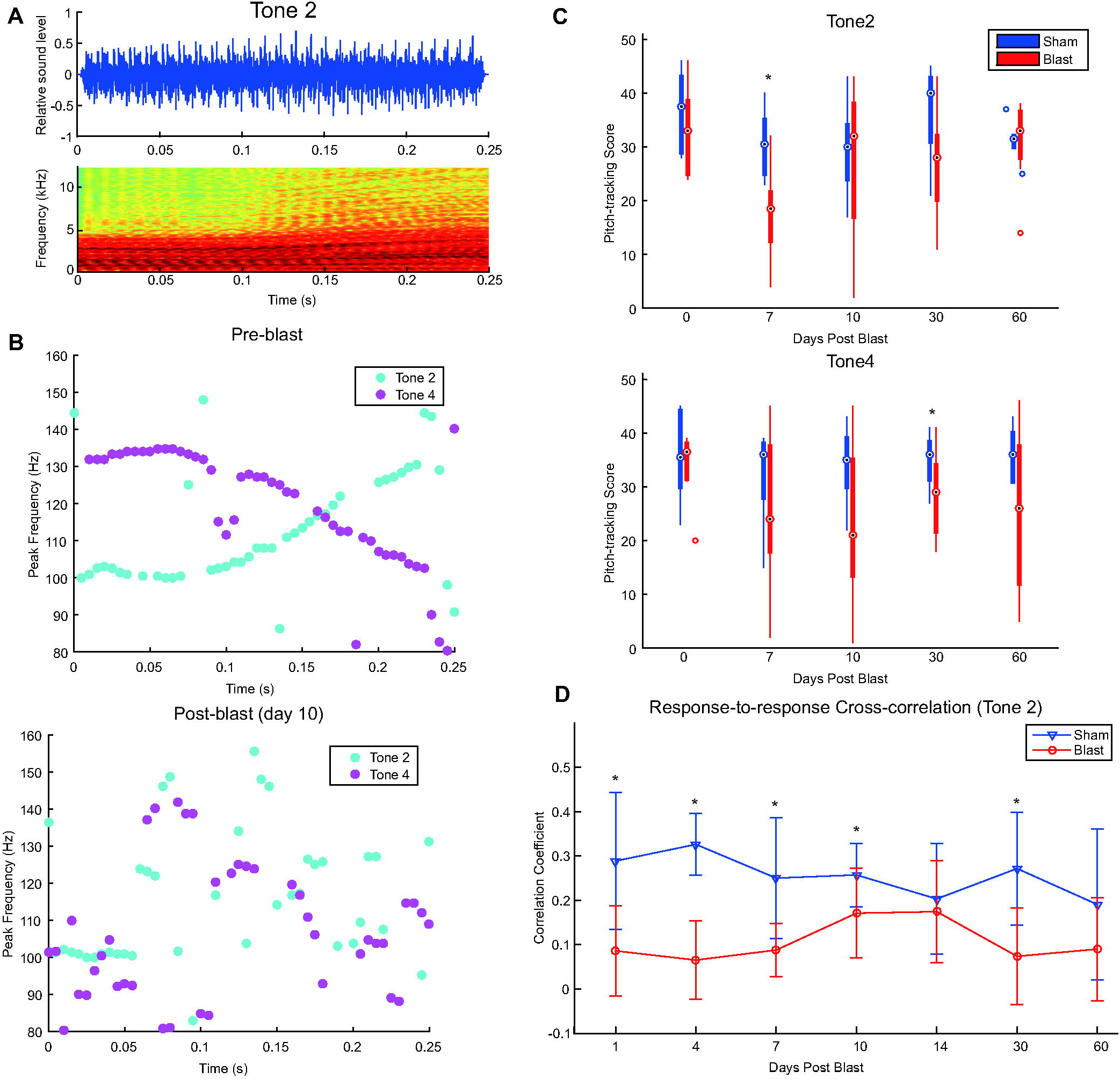
IRN Chinese Tone stimuli and responses from Blast (N=8) and Sham (N=6) rats during the first two months post-exposure at 30 dB above threshold, 32 iterations. A) Example waveform and spectrogram of IRN Tone 2 stimulus; B) Examples of Peak Frequency of IRN Evoked Potential in Pre-blast (score: Tone 2=36/51, Tone 4=39/51) and Post-blast Brain (day 10 post-blast, score: Tone 2=21/51, Tone 4=18/51) from an individual animal; C) Significant effect of Group (*) was seen in IRN Tone 2 (top, p=0.026) and Tone 4 (bottom, p=0.029) pitch-tracking score, though Simple Main Effect of is limited for individual time points; D) Cross-correlation of post-blast IRN responses to corresponding pre-blast responses. Significant differences (*) in correlation coefficients to pre-blast responses between Blast and Sham were observed in two waves: day 1-10, and day 30 (p<0.05, paired sign-rank).

For 8 kHz SAM, the effects of Noise level were applied differently between Groups, as significant interaction effects were observed between Noise and Group for 10 Hz (df=2.000, F=12.795, p=0.001, η^2^_p_=0.477) and 45 Hz (df=2.000, F=4.878, p=0.015, η^2^_p_=0.258) modulation frequencies, though not for 256 Hz (data not shown). Most notably, sham 8 kHz SAM EFRs showed greater resilience to competing noise at 10 Hz modulation frequency (Fig. 7C), contrary to the trends observed in nSAM. Modulation Depth affects FFT amplitude without regard to blast condition, with no significant interaction effects with Group observed. For 16 kHz SAM, none of the parameters tested had significantly different effects between Groups at 30 dB SL (not shown).

Overall, Blast and Sham animals generally decreased EFR amplitudes with increased noise, especially for 0 dB SNR. Similar to quiet, 45 Hz amplitudes were most affected, with increases in EFR amplitudes in Blast animals that were most pronounced in the 7-14 day window (Fig. 7B). The effects and interactions of blast exposure and competing noise were dependent on both modulation frequency and time after exposure..

### D. IRN

Time-varying IRN stimuli (Fig. 8A) were used to elicit frequency-following response (FFR) mimicking Mandarin tone 2 (T2, rising) and tone 4 (T4, falling) pitch contours to measure pitch-tracking ability using a broadband speech-like carrier at 30 dB SL, similar to what has been measured in human studies of auditory learning and hearing loss (Anderson et al. 2010, 2013; Skoe and Kraus 2010). We used autocorrelation interval contours that simulated pitches similar to the forms of rising (T2) and falling (T4) pitch contours of the Mandarin Chinese vowel /yi/ (Krishnan et al. 2014, 2015, 2017a, 2017b). IRN responses collected from channel 1 were evaluated based on the pitch-tracking score (Fig. 8B), which measures the number of time windows where the dominant autocorrelation frequency of the response matches that of the IRN stimulus autocorrelation frequency (see Methods). In general, we observed a loss of pitch-tracking fidelity in Blast animals over the two months post-exposure (Fig. 8B and 8C). Even for the most salient pitch (32 Iterations), blast exposure had a significant Group effect on pitch-tracking scores in both Tone 2 (df=1.000, F=6.495, p=0.026, η^2^_p_=0.351) and Tone 4 (df=1.000, F=6.115, p=0.029, η^2^_p_=0.338), with the largest mean differences on day 7-10. The interaction effect between Time and Group was not significant.

Blast exposure significantly changed the neural response’s morphology to IRN at 30 dB SL (p=0.016, paired sign-rank test, Fig. 8D), such that the cross-correlation between the pre-exposure response and the post-exposure response was much lower in the Blast group up to 30 days post-blast.

#### IRN iterations

As expected, reduced pitch salience, controlled by reducing iteration number, affected pitch-tracking responses in animals (df=4.000, F=41.697, p<0.001, η^2^_p_=0.777), also showing a significant interaction effect with Time post-exposure (df=20.000, F=1.722, p=0.031, η^2^_p_=0.125). Specifically, pitch-tracking performances to 32 iterations and 16 iterations worsened significantly up to 7-10 days post-exposure, with various degrees of recovery over the following time course. Both the Blast and Sham group exhibited worse pitch tracking with reduced iterations (salience) and to a similar degree. No significant interaction effects with Group were observed for Time and Iterations (Fig 9).

**Fig 9.**
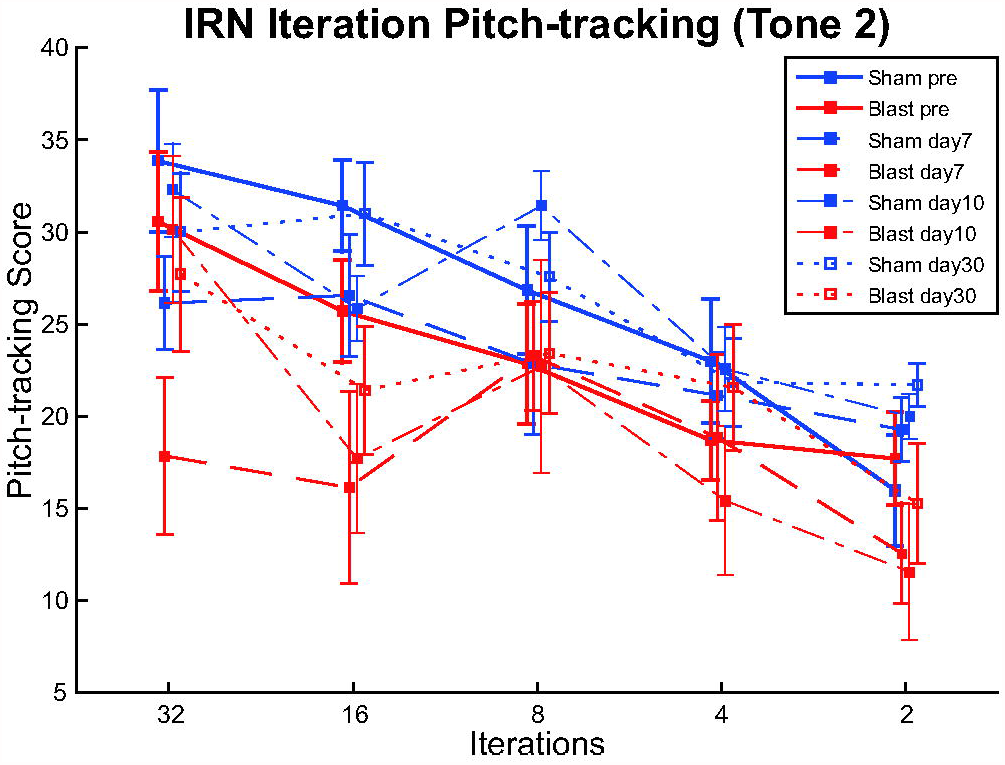
Pitch tracking scores of responses to IRN Tone 2 stimuli with pitch salience controlled by altering iteration number at different time points, at 30 dB above threshold. Though the effect of Iterations on pitch-tracking score was significant (p<0.001), no significant Iteration * Group interaction was observed.

## Discussion

This study examined the time course of recovery from a single mild blast injury using simple and complex auditory stimuli longitudinally at dense time points for two months. The largest blast-induced threshold shifts and changes in evoked potentials diminished within two weeks. At 30-60 days post-blast, lingering increases in click (but not tone) thresholds, decreases in MLR N1 (thlamocortical) amplitude, and declines in pitch-tracking of speech-like IRN pitch trajectories were observed. Compensating for threshold shift and using 30 dB sensation level for AM stimuli, we found that responses to sinusoidal AM stimuli in quiet or noise recovered within 14 days. The 7-14 day window was particularly rapid in the recovery of many auditory parameters.

### Lasting changes from a single mild blast

This study has examined injuries elicited by a single dorsal blast exposure with body shielding that did not result in tympanic membrane ruptures, which we and others characterize as a “mild” blast exposure. Therefore, the deficits observed may not be as drastic as that documented by some previous studies in which the injuries were caused by more intense or multiple exposures (Cho et al. 2013b; Du et al. 2013; Luo et al. 2014a, 2014b; Mahmood et al. 2014), often resulting in death or tympanic membrane rupture. The distribution of injuries also differed from models in which blast exposure comes from different orientations, as predicted in animals (Chavko et al. 2011; Dal Cengio Leonardi et al. 2012) and computational studies (Hua et al. 2017; Unnikrishnan et al. 2019). These differences in pressure wave amplitude, duration, and propagation patterns would affect both the distribution and severity of damage across the brain. Compared to other orientations, including top-facing exposure as in our model, head-facing exposure is known to produce the highest peak pressure and prolonged pressure wave propagation, while side-facing exposure produced lower peak pressure and pressure increase rate in rat model (Chavko et al. 2011; Dal Cengio Leonardi et al. 2012). Although these could change the potential mechanisms of recovery and compensation, it is likely that all blast exposures undergo a multi-stage recovery process similar to that observed in the present study. In our model, the overpressure blast wave passes through the entire rat brain, such that TBI can be observed throughout the brain, including the frontal cortex and in multiple thalamic regions (Walls et al. 2016), and it results in increased ventral BBB membrane permeation and inflammation, encompassing many subcortical auditory nuclei and axonal tracts. The non-invasive physiological measurements in this study may be indicators of more widespread blast damage in auditory and may be correlated with damage in non-auditory brain regions.

### ABR

We documented a >30 dB peak increase in threshold for click, 8 kHz, and 16 kHz (Fig. 2) during the first 4 days, consistent with the description of IHC and OHC disturbances across a wide range of frequencies due to blast overpressure as stated in multiple previous publications (Patterson and Hamernik 1997; Ewert et al. 2012; Race et al. 2017). Although this broadband threshold shift extended to the last time point at 60 days, the ∼10 dB difference would not be considered clinically relevant and suggests.

Rapid improvements in ABR threshold and wave amplitudes were observed in the 7-10 days recovery period for waves I, wave III, and wave V (Fig. 3). Notably, wave V amplitude recovered earlier than wave I, possibly indicating the role of compensation in auditory midbrain as one of the post-blast recovery mechanisms.

ABR parameters showed two waves of post-blast changes: one between 1-10 days post-exposure, and one 10-30 days, as evidenced by Figs. 2 and 3. We hypothesize that these two waves of deficits indicated a series of secondary biochemical impacts surrounding CAS (Laplaca et al. 1997; Knudsen and Øen 2003; Hamann et al. 2008; Garman et al. 2011; Säljö et al. 2011; Luo et al. 2014a, 2014b; Song et al. 2015; Walls et al. 2016). In the initial recovery window, we observed changes in ABR waveforms over and above those expected by threshold shifts, whereas for days 10 and afterwards, there were changes observed at 80 dB SPL but not for 30 dB SL. Our observations of blast recovery were mostly consistent with the notion of changes over the first week due to secondary damage that is substantially repaired over the second week.

### MLR

At 80 dB SPL, we found persistent deficits in thalamocortical and cortical transmission based on the N1, P2 and N2 peaks (Fig. 5A vs. B, C vs. D), which were affected at 30 and 60 days, even after the early P1 response had fully recovered (Fig 5E). These deficits were not present at 30 dB SL, suggesting that effects were at least partially due to small shifts in auditory thresholds (Fig. 5G). In veterans and the general population with lifetime noise exposure, MLR responses were shown to be smaller even when subjects had clinically normal audiograms, and there was some evidence of increased cortical gain (Valderrama et al. 2018; Bramhall et al. 2020). In another study with blast-exposed veterans, most of the changes in auditory-evoked potentials were correlated with hearing loss (Meehan et al. 2019). Together, these results suggest that hearing loss may be the main contributor to MLR changes leading to declines in suprathreshold responses.

### Amplitude Modulation EFRs

The current study extended an earlier study (Race et al. 2017) to include EFR responses to more challenging auditory stimuli, including lower modulation depth (Fig 6) and in the presence of modulated noise (Fig 7). The Race et al. (Race et al. 2017) study revealed differences in AM processing at 80 dB SPL between Blast and Sham animals, such that blast animals had lower AM FFR amplitudes mainly for AM frequencies ≤ 50 Hz. However, when the hearing threshold has been compensated, the differences in AM FFR amplitude diminished and even changed sign (Fig. 6), suggesting that both changes in audibility and changes in the gain of subcortical auditory system are critical contributors to AM FFR deficits in the blast-exposed auditory system. There are complicated interactions between the AM FFR amplitudes, blast exposure, and the presence of noise, evident as a persistent Group*Noise Level interaction effect in both nSAM and 8 kHz SAM. AM responses consist of contributions from multiple generators along the auditory neuraxis, with cortical generators contributing mainly to lower AMFs <50 Hz, and higher frequency AM responses limited to nuclei lower in the auditory neuraxis. The lack of blast-induced differences at higher AMFs distinguishes the blast-induced damage from age-related changes, which are most prominent at higher modulation frequencies (Parthasarathy et al. 2010, Parthasarathy and Bartlett 2012, Lai et al. 2017).

The differences in low-middle AMFs were manifested in opposed directions under slow (10 Hz) and middle (45 Hz) AMFs: notably, repeated measures testing showed that FFT amplitudes of 8 kHz SAM in noise are lower for Blast at 10 Hz modulation frequency (Day 4 quiet, 100% depth: Blast mean=0.496 mV, Sham mean=0.704 mV), but higher for Blast at 45 Hz (Day 4 quiet, 100% depth: Blast mean=0.898 mV, Sham mean=0.733 mV ; Fig 7C). This dichotomy is ripe for further study since the 10 Hz and 45 Hz modulations represent different temporal processing regimes and components of speech (Rosen 1992). If these modulation frequency bands are differentially altered by blast, it may skew the cochlear-filtered envelope and impair hearing in complex listening environments (Chabot-Leclerc et al. 2016).

### IRN EFRs

Complex temporal periodicity between 50 Hz and 500 Hz carries important speech information such as voicing, stress and intonation (Rosen 1992). The present study provided insights into blast-induced sound processing deficits through the use of an IRN stimulus that simulates Chinese intonations and whose pitch and salience can be reliably controlled, showing that IRN can be a useful diagnostic tool for neurotrauma. We found that even when click ABR thresholds have returned to subclinical threshold shifts, the deficits in pitch-tracking response to IRN tone stimuli, lingered at least 30 days post-exposure (Figs. 8D).

Both Blast and Sham animals showed an overall reduction in tracking with decreased salience through decreased iterations in IRN, but differential effects were noted mainly only in the first two weeks. A previous study showed that increased IRN iterations improved auditory stream segregation in normal hearing veterans more than hearing-impaired veterans (Thompson and Marozeau 2014). Our IRN data (Figs. 8-9) suggest that more dynamic and speech-like modulation changes do not recover quickly or completely from even a single mild blast exposure.

## Acknowledgments

The authors would like to thank Jonathan Tang, Brandon Coventry, Alex Sommers, Nanami Miyazaki, and all the other members in Central Auditory Processing Lab and Lab of Translational Neuroscience for their generous assistance in the completion of this study.

## Author contributions

Study concept and design: Edward Bartlett, Riyi Shi, Emily X. Han, Joseph M. Fernandez.

Animal Blast Exposure: Joseph M. Fernandez.

Electrophysiology: Emily X. Han, Caitlin Swanberg.

Data analysis and interpretation: Emily X. Han, Edward Bartlett, Joseph M. Fernandez.

Writing of the manuscript: Emily X. Han, Edward Bartlett.

Critical revision of the manuscript: Riyi Shi, Joseph M. Fernandez, Caitlin Swanberg. Study supervision and obtainment of funding: Edward Bartlett, Riyi Shi.

---

**Supplementary Table 1.**
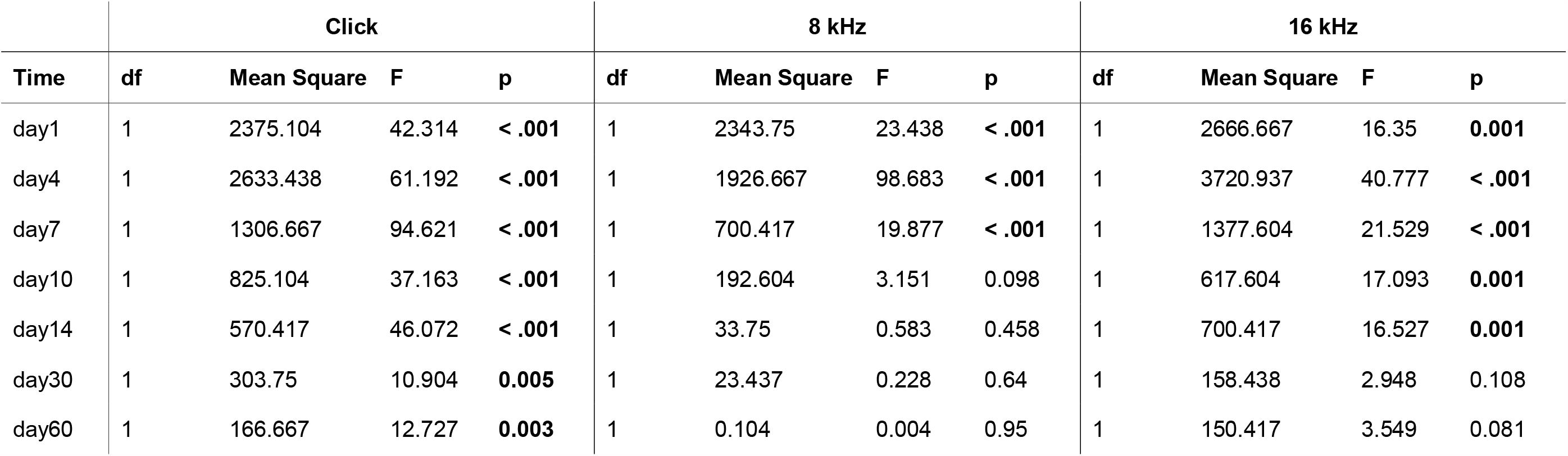
Simple main effects of Group on Click, 8 kHz and 16 kHz ABR threshold over time. Post-blast ABR thresholds of Blast (N=10) and Sham (N=6) groups are compared using rmANOVA at each time point recorded. A p<0.05 showed significant simple main effect of Group at that time point.

## References

Alsindi S, Patterson RD, Sayles M, Winter IM. The responses of single units to simple and complex sounds from the superior olivary complex of the Guinea pig. In: Acta Acustica united with Acustica. S. Hirzel Verlag GmbH, 2018, p. 856–859.

Alvarado JC, Fuentes-Santamaría V, Jareño-Flores T, Blanco JL, Juiz JM. Normal variations in the morphology of auditory brainstem response (ABR) waveforms: A study in wistar rats. Neurosci Res 73: 302–311, 2012.

Anderson S, Parbery-Clark A, White-Schwoch T, Kraus N. Auditory brainstem response to complex sounds predicts self-reported speech-in-noise performance. J Speech, Lang Hear Res 56: 31–43, 2013.

Anderson S, Skoe E, Chandrasekaran B, Kraus N. Neural timing is linked to speech perception in noise. J Neurosci 30: 4922–4926, 2010.

Barth DS, Shi Di. The functional anatomy of middle latency auditory evoked potentials. Brain Res 565: 109–115, 1991.

Bendor D, Wang X. The neuronal representation of pitch in primate auditory cortex. Nature 436: 1161–1165, 2005.

Berger G, Finkelstein Y, Avraham S, Himmelfarb M. Patterns of hearing loss in non-explosive blast injury of the ear. J Laryngol Otol 111: 1137–1141, 1997.

Bharadwaj HM, Masud S, Mehraei G, Verhulst S, Shinn-Cunningham BG. Individual differences reveal correlates of hidden hearing deficits. J Neurosci 35: 2161–2172, 2015.

Bramhall NF, Niemczak CE, Kampel SD, Billings CJ, McMillan GP. Evoked Potentials Reveal Noise Exposure–Related Central Auditory Changes Despite Normal Audiograms. Am J Audiol 29: 152–164, 2020.

Bressler S, Goldberg H, Shinn-Cunningham B. Sensory coding and cognitive processing of sound in Veterans with blast exposure. Hear Res 349: 98–110, 2017.

Brett B, Krishnan G, Barth DS. The effects of subcortical lesions on evoked potentials and spontaneous high frequency (gamma-band) oscillating potentials in rat auditory cortex. Brain Res 721: 155–166, 1996.

Caspary DM, Ling L, Turner JG, Hughes LF. Inhibitory neurotransmission, plasticity and aging in the mammalian central auditory system. J Exp Biol 211: 1781–1791, 2008.

Caspary DM, Schatteman T a, Hughes LF. Age-related changes in the inhibitory response properties of dorsal cochlear nucleus output neurons: role of inhibitory inputs. J Neurosci 25: 10952–9, 2005.

Cave K, Cornish EM, Chandler DW. Blast Injury of the Ear: Clinical Update from the Global War on Terror. Mil Med 172: 726–730, 2007.

Chabot-Leclerc A, MacDonald EN, Dau T. Predicting binaural speech intelligibility using the signal-to-noise ratio in the envelope power spectrum domain. J Acoust Soc Am 140: 192–205, 2016.

Chavko M, Watanabe T, Adeeb S, Lankasky J, Ahlers ST, McCarron RM. Relationship between orientation to a blast and pressure wave propagation inside the rat brain. J Neurosci Methods 195: 61–66, 2011.

Cho HJ, Sajja VSSS, VandeVord PJ, Lee YW. Blast induces oxidative stress, inflammation, neuronal loss and subsequent short-term memory impairment in rats. Neuroscience 253: 9–20, 2013a.

Cho S Il, Gao SS, Xia A, Wang R, Salles FT, Raphael PD, Abaya H, Wachtel J, Baek J, Jacobs D, Rasband MN, Oghalai JS. Mechanisms of Hearing Loss after Blast Injury to the Ear. PLoS One 8, 2013b.

Cohen JT, Ziv G, Bloom J, Zikk D, Rapoport Y, Himmelfarb MZ. Blast injury of the ear in a confined space explosion: Auditory and vestibular evaluation. Isr Med Assoc J 4: 559–562, 2002.

Dal Cengio Leonardi A, Keane NJ, Bir CA, Ryan AG, Xu L, VandeVord PJ. Head orientation affects the intracranial pressure response resulting from shock wave loading in the rat. J Biomech 45: 2595–2602, 2012.

Deiber MP, Ibañez V, Fischer C, Perrin F, Mauguière F. Sequential mapping favours the hypothesis of distinct generators for Na and Pa middle latency auditory evoked potentials. Electroencephalogr Clin Neurophysiol Evoked Potentials 71: 187–197, 1988.

DePalma RG, Burris DG, Champion HR, Hodgson MJ. Current concepts: Blast injuries. N. Engl. J. Med. 352Massachusetts Medical Society: 1335–1342, 2005.

Di S, Barth DS. The functional anatomy of middle-latency auditory evoked potentials: Thalamocortical connections. J Neurophysiol 68: 425–431, 1992.

Du X, Ewert DL, Cheng W, West MB, Lu J, Li W, Floyd RA, Kopke RD. Effects of antioxidant treatment on blast-induced brain injury. PLoS One 8: e80138, 2013.

Ewert DL, Lu J, Li W, Du X, Floyd R, Kopke R. Antioxidant treatment reduces blast-induced cochlear damage and hearing loss. Hear Res 285: 29–39, 2012.

Gallun FJ, Diedesch AC, Kubli LR, Walden TC, Folmer RL, Samantha Lewis S, McDermott DJ, Fausti SA, Leek MR. Performance on tests of central auditory processing by individuals exposed to high-intensity blasts. J Rehabil Res Dev 49: 1005–1024, 2012a.

Gallun FJ, Lewis MS, Folmer RL, Diedesch AC, Aud ;, Kubli LR, McDermott DJ, Walden TC, Fausti S a, Lew HL, Leek MR. Implications of blast exposure for central auditory function: a review. J Rehabil Res Dev 49: 1059–74, 2012b.

Garman RH, Jenkins LW, Switzer RC, Bauman RA, Tong LC, Swauger P V., Parks SA, Ritzel D V., Dixon CE, Clark RSB, Bayir H, Kagan V, Jackson EK, Kochanek PM. Blast Exposure in Rats with Body Shielding Is Characterized Primarily by Diffuse Axonal Injury. J Neurotrauma 28: 947–959, 2011.

Greenhouse SW, Geisser S. On methods in the analysis of profile data. Psychometrika 24: 95–112, 1959.

Van Haesendonck G, Van Rompaey V, Gilles A, Topsakal V, Van de Heyning P. Otologic Outcomes After Blast Injury: The Brussels Bombing Experience. Otol Neurotol 39: 1250–1255, 2018.

Hamann K, Durkes A, Ouyang H, Uchida K, Pond A, Shi R. Critical role of acrolein in secondary injury following ex vivo spinal cord trauma. J Neurochem 107: 712–721, 2008.

Hickox AE, Liberman MC. Is noise-induced cochlear neuropathy key to the generation of hyperacusis or tinnitus? J Neurophysiol 111: 552–564, 2014.

Hua Y, Wang Y, Gu L. Primary blast waves induced brain dynamics influenced by head orientations. Biomed Eng Lett 7: 253–259, 2017.

Joris PX, Schreiner CE, Rees A. Neural Processing of Amplitude-Modulated Sounds. American Physiological Society, 2004.

Kerr AG. The effects of blast on the ear. J Laryngol Otol 94: 107–110, 1980.

Knudsen SK, Øen EO. Blast-induced neurotrauma in whales. Neurosci Res 46: 377–386, 2003.

Krishnan A, Gandour JT, Suresh CH. Cortical pitch response components show differential sensitivity to native and nonnative pitch contours. Brain Lang 138: 51–60, 2014.

Krishnan A, Gandour JT, Suresh CH. Pitch processing of dynamic lexical tones in the auditory cortex is influenced by sensory and extrasensory processes. Eur J Neurosci 41: 1496–1504, 2015.

Krishnan A, Gandour JT, Xu Y, Suresh CH. Language-dependent changes in pitch-relevant neural activity in the auditory cortex reflect differential weighting of temporal attributes of pitch contours. J Neurolinguistics 41: 38–49, 2017a.

Krishnan A, Suresh CH, Gandour JT. Changes in pitch height elicit both language-universal and language-dependent changes in neural representation of pitch in the brainstem and auditory cortex. Neuroscience 346: 52–63, 2017b.

Kubli LR, Brungart D, Northern J. Effect of Blast Injury on Auditory Localization in Military Service Members. Ear Hear 39: 457–469, 2018.

Kujawa SG, Liberman MC. Synaptopathy in the noise-exposed and aging cochlea: Primary neural degeneration in acquired sensorineural hearing loss. Hear Res 330: 191–199, 2015.

Lai J, Bartlett EL. Age-related shifts in distortion product otoacoustic emissions peak-ratios and amplitude modulation spectra. Hear Res 327: 186–198, 2015.

Lai J, Bartlett EL. Masking Differentially Affects Envelope-following Responses in Young and Aged Animals. Neuroscience 386: 150–165, 2018.

Lai J, Sommer AL, Bartlett EL. Age-related changes in envelope-following responses at equalized peripheral or central activation. Neurobiol Aging 58: 191–200, 2017.

Laplaca MC, Lee VMY, Thibault LE. An in vitro model of traumatic neuronal injury: Loading rate-dependent changes in acute cytosolic calcium and lactate dehydrogenase release. J Neurotrauma 14: 355–368, 1997.

Leung LY, VandeVord PJ, Dal Cengio AL, Bir C, Yang KH, King AI. Blast related neurotrauma: A review of cellular injury. MCB Mol Cell Biomech 5: 155–168, 2008.

Lew HL, Garvert DW, Pogoda TK, Hsu P Te, Devine JM, White DK, Myers PJ, Goodrich GL. Auditory and visual impairments in patients with blast-related traumatic brain injury: Effect of dual sensory impairment on functional independence measure. J Rehabil Res Dev 46: 819–826, 2009.

Liberman MC. Noise-induced and age-related hearing loss: New perspectives and potential therapies. F1000Research 6: 1–11, 2017.

Liégeois-Chauvel C, Musolino A, Badier JM, Marquis P, Chauvel P. Evoked potentials recorded from the auditory cortex in man: evaluation and topography of the middle latency components. Electroencephalogr Clin Neurophysiol Evoked Potentials 92: 204–214, 1994.

Luo H, Pace E, Zhang X, Zhang J. Blast-Induced tinnitus and spontaneous firing changes in the rat dorsal cochlear nucleus. J Neurosci Res 92: 1466–1477, 2014a.

Luo H, Pace E, Zhang X, Zhang J. Blast-induced tinnitus and spontaneous activity changes in the rat inferior colliculus. Neurosci Lett 580: 47–51, 2014b.

Mahmood G, Mei Z, Hojjat H, Pace E, Kallakuri S, Zhang JS. Therapeutic effect of sildenafil on blast-induced tinnitus and auditory impairment. Neuroscience 269: 367–382, 2014.

Mao JC, Pace E, Pierozynski P, Kou Z, Shen Y, Vandevord P, Haacke EM, Zhang X, Zhang J. Blast-induced tinnitus and hearing loss in rats: Behavioral and imaging assays. J Neurotrauma 29: 430–444, 2012.

Masri S, Zhang LS, Luo H, Pace E, Zhang J, Bao S. Blast Exposure Disrupts the Tonotopic Frequency Map in the Primary Auditory Cortex. Neuroscience 379: 428–434, 2018.

McGee T, Kraus N. Auditory development reflected by middle latency response. Ear Hear 17: 419–429, 1996.

McGee T, Kraus N, Comperatore C, Nicol T. Subcortical and cortical components of the MLR generating system. Brain Res 544: 211–220, 1991.

McGee T, Kraus N, Littman T, Nicol T. Contributions of medial geniculate body subdivisions to the middle latency response. Hear Res 61: 147–154, 1992.

Meehan A, Hebert D, Deru K, Weaver LK. Longitudinal study of hyperbaric oxygen intervention on balance and affective symptoms in military service members with persistent post-concussive symptoms. J Vestib Res Equilib Orientat 29: 205–219, 2019.

Musiek F, Nagle S. The middle latency response: A review of findings in various central nervous system lesions. J. Am. Acad. Audiol. 29American Academy of Audiology: 855–867, 2018.

Parthasarathy A, Bartlett E. Two-channel recording of auditory-evoked potentials to detect age-related deficits in temporal processing. Hear Res 289: 52–62, 2012.

Parthasarathy A, Bartlett EL. Age-related auditory deficits in temporal processing in F-344 rats. Neuroscience 192: 619–630, 2011.

Parthasarathy A, Cunningham PA, Bartlett EL. Age-related differences in auditory processing as assessed by amplitude-modulation following responses in quiet and in noise. Front Aging Neurosci 2: 3389, 2010.

Parthasarathy A, Datta J, Torres JAL, Hopkins C, Bartlett EL. Age-related changes in the relationship between auditory brainstem responses and envelope-following responses. JARO - J Assoc Res Otolaryngol 15: 649–661, 2014.

Parthasarathy A, Kujawa SG. Synaptopathy in the aging cochlea: Characterizing early-neural deficits in auditory temporal envelope processing. J Neurosci 38: 7108–7119, 2018.

Patterson JH, Hamernik RP. Blast overpressure induced structural and functional changes in the auditory system. 1997.

Peter V, Wong K, Narne VK, Sharma M, Purdy SC, McMahon C. Assessing spectral and temporal processing in children and adults using Temporal Modulation Transfer Function (TMTF), Iterated Ripple Noise (IRN) perception, and Spectral Ripple Discrimination (SRD). J Am Acad Audiol 25: 210–218, 2014.

Phillips DJ, Schei JL, Meighan PC, Rector DM. State-Dependent Changes in Cortical Gain Control as Measured by Auditory Evoked Responses to Varying Intensity Stimuli. Sleep 34: 1527–1537, 2011.

Plack CJ, Barker D, Prendergast G. Perceptual consequences of “hidden” hearing loss. Trends Hear 18: 233121651455062, 2014.

Rabang CF, Parthasarathy A, Venkataraman Y, Fisher ZL, Gardner SM, Bartlett EL. A computational model of inferior colliculus responses to amplitude modulated sounds in young and aged rats. Front Neural Circuits 6: 77, 2012.

Race N, Lai J, Shi R, Bartlett EL. Differences in postinjury auditory system pathophysiology after mild blast and nonblast acute acoustic trauma. J Neurophysiol 118: 782–799, 2017.

Rafaels K, “Dale” Bass CR, Salzar RS, Panzer MB, Woods W, Feldman S, Cummings T, Capehart B. Survival Risk Assessment for Primary Blast Exposures to the Head. J Neurotrauma 28: 2319–2328, 2011.

Remenschneider AK, Lookabaugh S, Aliphas A, Brodsky JR, Devaiah AK, Dagher W, Grundfast KM, Heman-Ackah SE, Rubin S, Sillman J, Tsai AC, Vecchiotti M, Kujawa SG, Lee DJ, Quesnel AM. Otologic outcomes after blast injury: The Boston Marathon experience. Otol Neurotol 35: 1825–1834, 2014.

Ritenour AE, Wickley A, Ritenour JS, Kriete BR, Blackbourne LH, Holcomb JB, Wade CE. Tympanic membrane perforation and hearing loss from blast overpressure in Operation Enduring Freedom and Operation Iraqi Freedom wounded. J Trauma 64, 2008.

Rosen S. Temporal information in speech: acoustic, auditory and linguistic aspects. Philos. Trans. R. Soc. Lond. B. Biol. Sci. 336The Royal Society London: 367–373, 1992.

Säljö A, Mayorga M, Bolouri H, Svensson B, Hamberger A. Mechanisms and pathophysiology of the low-level blast brain injury in animal models. Neuroimage 54: S83–S88, 2011.

Saunders GH, Frederick MT, Arnold M, Silverman S, Chisolm TH, Myers P. Auditory difficulties in blast-exposed veterans with clinically normal hearing. J Rehabil Res Dev 52: 343–360, 2015.

Simpson G V., Knight RT, Brailowsky S, Prospero-Garcia O, Scabini D. Altered peripheral and brainstem auditory function in aged rats. Brain Res 348: 28–35, 1985.

Simpson MIG, Prendergast G. Auditory magnetic evoked responses. In: Handbook of Clinical Neurophysiology. 2013, p. 253–270.

Skoe E, Kraus N. Auditory Brain Stem Response to Complex Sounds: A Tutorial. Ear Hear 31: 302–324, 2010.

Song S, Race NS, Kim A, Zhang T, Shi R, Ziaie B. A Wireless Intracranial Brain Deformation Sensing System for Blast-Induced Traumatic Brain Injury. Sci Rep 5: 1–10, 2015.

Šuta D, Rybalko N, Pelánová J, PopelářJ, Syka J. Age-related changes in auditory temporal processing in the rat. Exp Gerontol 46: 739–746, 2011.

Swaminathan J, Krishnan A, Gandour JT. Pitch encoding in speech and nonspeech contexts in the human auditory brainstem. Neuroreport 19: 1163–1167, 2008.

Thompson W, Marozeau J. Auditory Stream Segregation: Boundaries. In: Music in the Social and Behavioral Sciences: An Encyclopedia. 2014.

Tichko P, Skoe E. Frequency-dependent fine structure in the frequency-following response: The byproduct of multiple generators. Hear Res 348: 1–15, 2017.

Tukey JW. Comparing Individual Means in the Analysis of Variance. Biometrics 5: 99, 1949.

Unnikrishnan G, Mao H, Sundaramurthy A, Bell ED, Yeoh S, Monson K, Reifman J. A 3-D Rat Brain Model for Blast-Wave Exposure: Effects of Brain Vasculature and Material Properties. Ann Biomed Eng 47: 2033–2044, 2019.

Valderrama JT, Beach EF, Yeend I, Sharma M, Van Dun B, Dillon H. Effects of lifetime noise exposure on the middle-age human auditory brainstem response, tinnitus and speech-in-noise intelligibility. Hear Res 365: 36–48, 2018.

Viana LM, O’Malley JT, Burgess BJ, Jones DD, Oliveira CACP, Santos F, Merchant SN, Liberman LD, Liberman MC. Cochlear neuropathy in human presbycusis: Confocal analysis of hidden hearing loss in post-mortem tissue. Hear Res 327: 78–88, 2015.

Wagner L, Plontke SK, Rahne T. Perception of Iterated Rippled Noise Periodicity in Cochlear Implant Users. Audiol Neurotol 22: 104–115, 2017.

Walls MK, Race N, Zheng L, Vega-Alvarez SM, Acosta G, Park J, Shi R. Structural and biochemical abnormalities in the absence of acute deficits in mild primary blast-induced head trauma. J Neurosurg 124: 675–686, 2016.

Walton JP. Timing is everything: Temporal processing deficits in the aged auditory brainstem. Hear Res 264: 63–69, 2010.

Wang H, Turner JG, Ling L, Parrish JL, Hughes LF, Caspary DM. Age-related changes in glycine receptor subunit composition and binding in dorsal cochlear nucleus. Neuroscience 160: 227–239, 2009.

Wang X, Lu T, Bendor D, Bartlett E. Neural coding of temporal information in auditory thalamus and cortex. Neuroscience 154: 294–303, 2008.

Yost WA. Pitch of iterated rippled noise. J Acoust Soc Am 100: 511–518, 1996a.

Yost WA. Pitch strength of iterated rippled noise. J Acoust Soc Am 100: 3329–3335, 1996b.

