## Supplemental Table 1 for "Longitudinal Auditory Pathophysiology Following Mild Blast Induced Trauma"

**Supplementary Table 1**

|  | **Click** | | | | **8 kHz** | | | | **16 kHz** | | | |
| --- | --- | --- | --- | --- | --- | --- | --- | --- | --- | --- | --- | --- |
| **Time** | **df** | **Mean Square** | **F** | **p** | **df** | **Mean Square** | **F** | **p** | **df** | **Mean Square** | **F** | **p** |
| day1 | 1 | 2375.104 | 42.314 | **< .001** | 1 | 2343.75 | 23.438 | **< .001** | 1 | 2666.667 | 16.35 | **0.001** |
| day4 | 1 | 2633.438 | 61.192 | **< .001** | 1 | 1926.667 | 98.683 | **< .001** | 1 | 3720.937 | 40.777 | **< .001** |
| day7 | 1 | 1306.667 | 94.621 | **< .001** | 1 | 700.417 | 19.877 | **< .001** | 1 | 1377.604 | 21.529 | **< .001** |
| day10 | 1 | 825.104 | 37.163 | **< .001** | 1 | 192.604 | 3.151 | 0.098 | 1 | 617.604 | 17.093 | **0.001** |
| day14 | 1 | 570.417 | 46.072 | **< .001** | 1 | 33.75 | 0.583 | 0.458 | 1 | 700.417 | 16.527 | **0.001** |
| day30 | 1 | 303.75 | 10.904 | **0.005** | 1 | 23.437 | 0.228 | 0.64 | 1 | 158.438 | 2.948 | 0.108 |
| day60 | 1 | 166.667 | 12.727 | **0.003** | 1 | 0.104 | 0.004 | 0.95 | 1 | 150.417 | 3.549 | 0.081 |

**Supplementary Table 1. Simple main effects of Group on Click, 8 kHz and 16 kHz ABR threshold over time.** Post-blast ABR thresholds of Blast (N=10) and Sham (N=6) groups are compared using rmANOVA at each time point recorded. A p<0.05 showed significant simple main effect of Group at that time point.
